# Mapping targetable sites on the human surfaceome for the design of novel binders

**DOI:** 10.1101/2024.12.16.628626

**Authors:** Petra E. M. Balbi, Ahmed Sadek, Anthony Marchand, Ta-Yi Yu, Jovan Damjanovic, Sandrine Georgeon, Joseph Schmidt, Simone Fulle, Che Yang, Hamed Khakzad, Bruno E. Correia

## Abstract

The human cell surfaceome, integral to cell communication and disease mechanisms, presents a prime target for therapeutic intervention. *De novo* protein binder design against these cell surface proteins offers a promising yet underexplored strategy for drug development. However, the vast search space and limited data on natural or competitive binders have historically limited experimental success. In this study, we systematically analyzed the entire human surfaceome, identifying approximately 4,500 targetable sites and introducing high-quality binder seeds tailored for protein design applications. To validate these seeds, we implemented two experimental approaches (protein scaffolding and peptide cyclization) on three representative targets (FGFR2, IFNAR2, and HER3). Our results revealed a high success rate, emphasizing the precision and therapeutic potential of these seeds, as well as the need for constant improvements of computational protein design pipelines utilizing machine learning and physics-based methods. Additionally, we present SURFACE-Bind, an interactive database offering open access to all generated data. The high-throughput computational design methods and target-specific binder seeds established here pave the way for a new generation of targeted therapeutics for the human surfaceome.

## Introduction

The diverse array of human cell surface proteins, known as the human surfaceome [1], plays a crucial role in cell-to-cell communication and the transmission of signals from the outside to the inside of cells. Because these surface proteins are easily accessible, the surfaceome represents a rich resource for drug targeting and has emerged as a promising avenue for therapeutic development. It is estimated that the human surfaceome includes around 2,886 proteins, categorized into different families and subfamilies [2]. While some of these proteins, such as programmed death-1 (PD-1) [3], human epidermal growth factor receptor 2 (HER2) [4], and epidermal growth factor receptor (EGFR) [5], have been extensively targeted in protein design efforts, many others remain poorly understood or unexplored.

Computational design of *de novo* protein binders from scratch poses several challenges and requires innovative strategies to unravel the complexities of molecular interactions. Recent advances in deep learning have significantly enhanced protein design approaches [6], allowing for very precise structural accuracy and higher experimental success rates [7], [8], [9]. However, this remains a difficult task, particularly in cases where there are multiple potential binding sites on a target protein and limited or no available data on natural protein binders. To address these challenges, we recently introduced MaSIF-seed-search [10], a geometric deep-learning framework designed specifically for protein surfaces. This framework generates “fingerprints” that capture both the geometric and chemical features essential for protein–protein interactions (PPIs). Our findings demonstrated that these fingerprints provide insights into the structural complementarity of molecular complexes and also present new possibilities for designing novel protein binders.

Building on the significance of the human surfaceome and the success of the MaSIF framework, we have now systematically evaluated all proteins within the human surfaceome, identifying approximately 4,500 potential binding sites across 2,886 proteins, which could be leveraged in future protein design studies. We further assessed the binding propensity of each site based on a combination of geometric and chemical scores in its unbound state. These predictions assess the feasibility of targeting each site through pairwise docking of 640,000 seed structures. This analysis enabled us to propose a set of high-quality binder seeds together with seed-bound-state scores that can be used to guide and advance future protein design efforts. By establishing a comprehensive database of these high-quality seeds targeting every binding site in the human surfaceome, our approach significantly enhances the scale and efficiency of peptide and protein design efforts, paving the way for new opportunities in developing therapeutics. All data generated in this study are freely accessible through the interactive SURFACE-Bind database (https://surface-bind.inria.fr/), enabling users to explore target proteins of interest and to further design protein binders by utilizing the detected seeds.

## Results

### Computational analysis of the human surfaceome

We performed a large-scale comprehensive analysis of the targetability of the human surfaceome. As indicated in **Figure 1A**, we targeted all proteins from the human *in silico* surfaceome database, SURFY, which encompasses 2,886 proteins that are either experimentally determined or predicted to be surface proteins [2]. These proteins are classified functionally into six main families: receptors, transporters, enzymes, miscellaneous, unclassified, and unmatched. For each protein entry, the corresponding AlphaFold2 (AF2) [11] model was obtained and filtered based on the predicted local distance difference test (pLDDT) (see Methods section). A large-scale site prediction approach was carried out via the molecular surface interaction fingerprints (MaSIF-site) framework [12], mapping the surface of each protein into a point cloud and predicting which surface points are more likely to interact with other proteins. To define the binding sites and to distinguish between the boundaries of two binding sites in close proximity, these points were subsequently clustered using density-based spatial clustering of applications with noise (DBSCAN). As an end result, we predicted a total number of 4,500 target sites excluding the transmembrane domains. The average number of sites per family and sub-family is shown in **Figure 1B**, where the GPCR and SLC obtained the lowest number of sites, while Kinase and Other Receptors have an average of 6 binding sites per protein. The frequency of detected sites per family and sub-family is depicted in **Figure 1C**.

**Figure 1:**
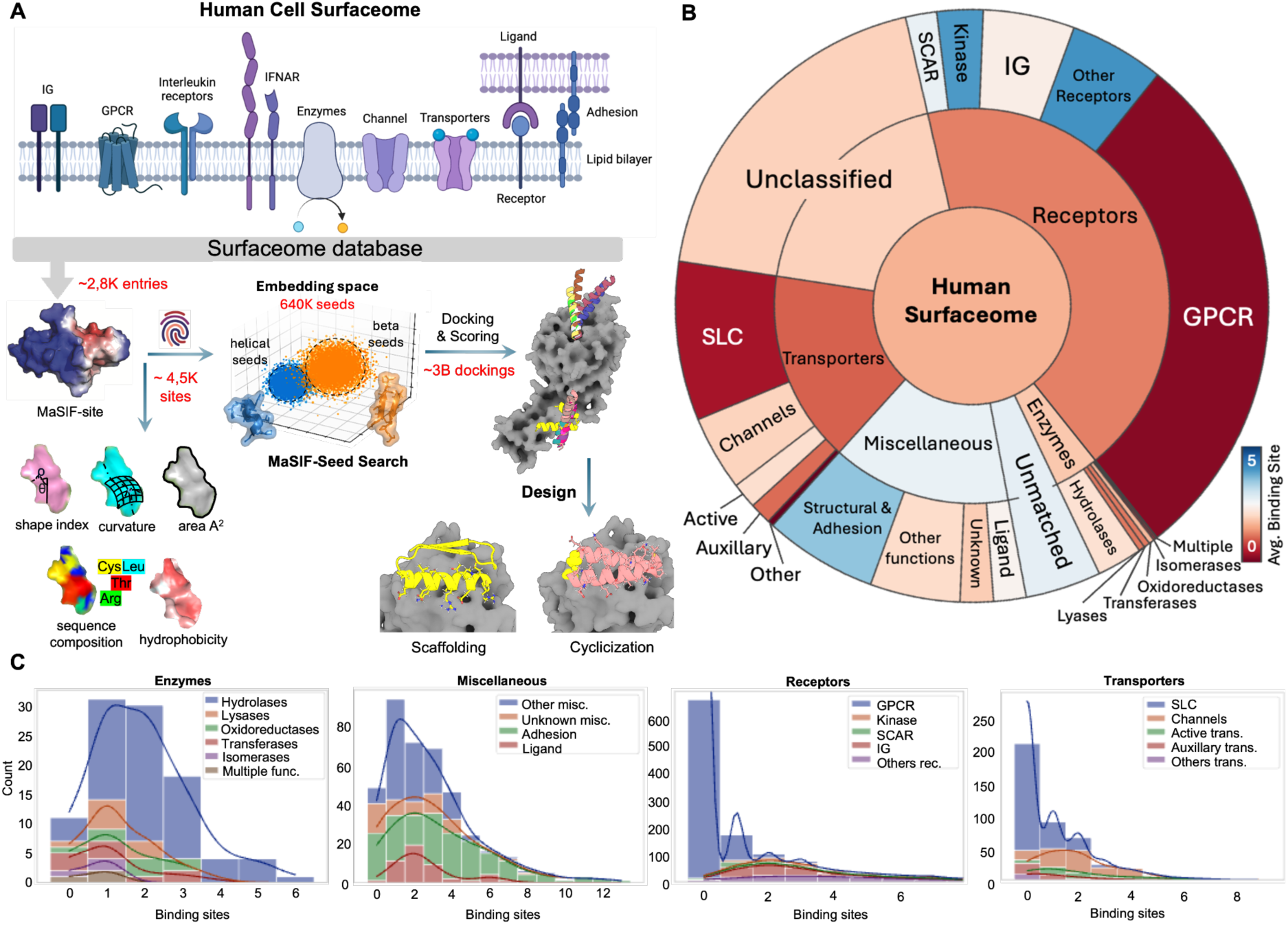
Overview of computational workflow and targetable sites in the human surfaceome. **A.** Schematic overview of the computational workflow. The Surfaceome structures were gathered from the SURFY database, comprising 2,886 entries [2]. High propensity protein binding sites were predicted using MaSIF-site [12], resulting in a total of 4,500 binding sites. These binding sites were then targeted by a library of 640,000 protein fragments based on the MaSIF-seed algorithm [10], and followed by pairwise docking with the evaluation of around 3 billion interactions. Subsequently, two protein design approaches, including scaffolding and cyclic peptide design, were then applied on representative proteins to evaluate the quality of seeds under experimental settings. **B.** Average predicted binding sites per family. Segment size corresponds to the number of proteins in each family, with GPCRs having the largest protein count but the fewest predicted binding sites, primarily located within the transmembrane (TM) domain. **C.** Binding site frequency on the four main protein families. Each sub-family is shown in a different color. Miscellaneous and transporter main classes have several members with multiple predicted binding interfaces, where some members in adhesion proteins contain more than 12 binding interfaces.

To evaluate the targetability of the predicted sites, we computed two sets of scores, including unbound- and seed-bound-state scores. The unbound scores is composed of geometric and chemical features such as shape index, curvature (concave, convex, flat), estimated binding interface area ( A^2^), sequence composition, solvent accessible surface area (SASA), and the hydrophobicity scale [13]. The distribution of unbound-state features across each family and sub-family are shown in **Supplementary Figure S1**, and **S2**. Analyzing the sequence composition of the predicted sites, we noted that leucine, valine and serine are the most frequent residues in the predicted binding sites (**Figure 2A**), in line with the typical hydrophobic character found in PPI sites.

**Figure 2:**
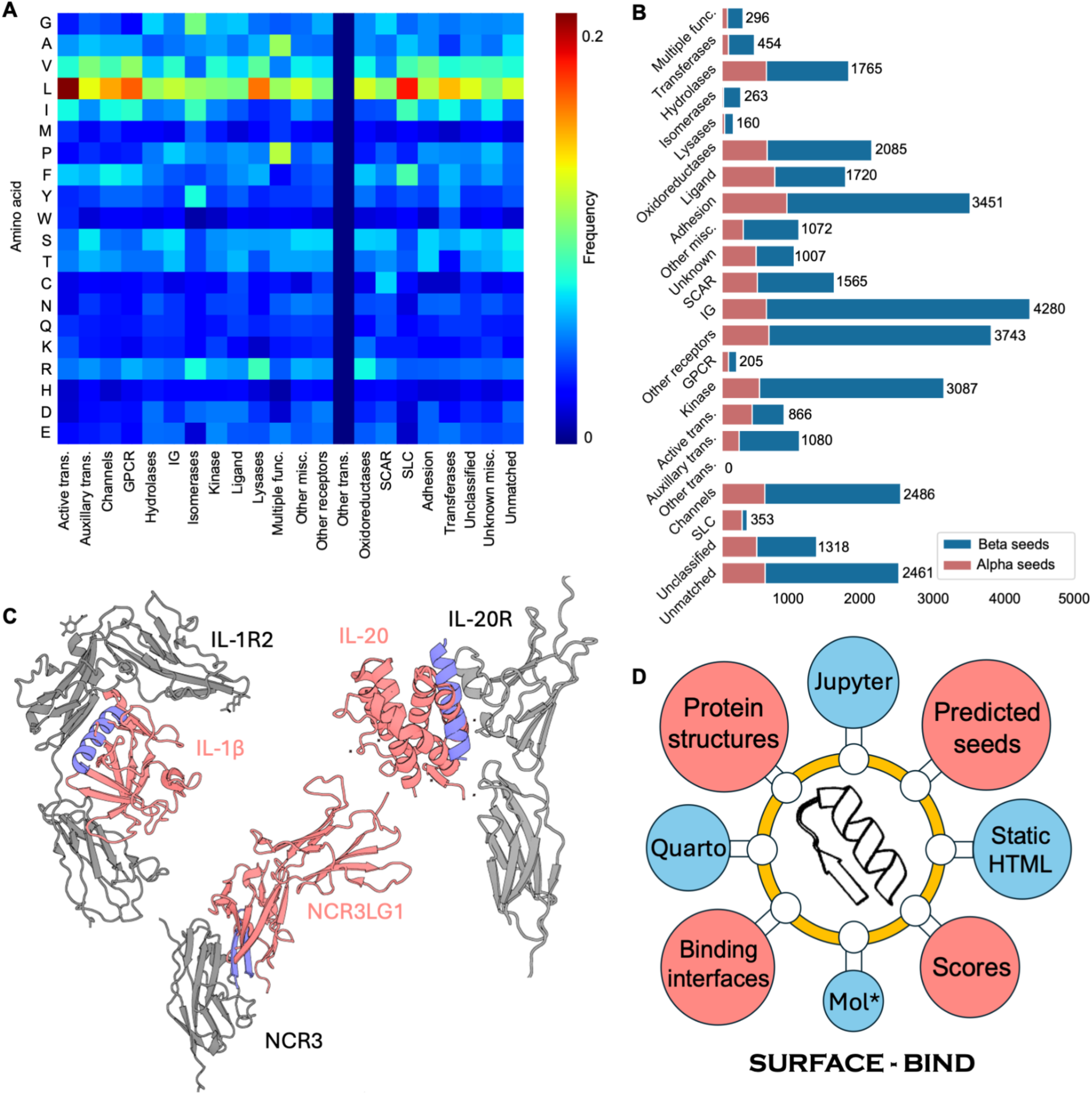
Mapping of the targetable human surfaceome. **A.** The amino acid frequency on the predicted binding sites for all sub-families. Leucine is the most frequent amino acid found in the predicted binding sites across all main classes and specifically in the transporters family. Valine and serine are the second and third frequent amino acids, respectively. **B.** Average number of alpha and beta seeds per sub-family. The number of detected alpha seeds is lower than the number of beta seeds, primarily due to the difference in the initial size of their databases. Although families with a higher number of binding sites obtained a higher number of seeds, the Ig subfamily showed a significant bind-ability to beta seeds. **C.** Examples include entries with natural binders (gray and red) and detected inhibitory seeds (blue) that disrupt the interaction. **D.** The SURFACE-Bind database provides the data openly available through static and interactive HTML pages.

Next, in order to calculate the bound-state scores, we targeted these sites by a library of 640,000 small helical and beta strand fragments (seeds) derived from the Protein Data Bank (PDB). We performed and evaluated nearly 3 billion pairwise dockings resulting in bound-state scores and a selection of high-quality seeds for each target site with highest complementarity matching score [10] (depicted in **Figure 2B**, **Supplementary Figure S1**). On average, receptors within the Ig family exhibit the highest number of matching seeds. As depicted by three examples, the detected PPI sites have also been found to overlap with the natural ligand binding site (**Figure 2C**), which could potentially disrupt the biological function of the protein. During the seed matching process, Masif-seed was also capable of identifying diverse structural seeds for docking a given predicted PPI site, thereby providing an opportunity for subsequent binder generation.

In order to make the data freely available to the community, we created a novel database called SURFACE-Bind (**Figure 2D**). This database comprises static HTML pages for individual entries, each associated with a specific UniProt ID. Each page contains detailed information on the protein structure, predicted binding interfaces, detected seeds, and several visual plots showcasing the distribution of chemical properties for both the binding interfaces and the corresponding seeds. All associated files can be conveniently downloaded in zip format at the bottom of each entry page, providing easy access to specific data.

### *De novo* designed binders against cell surface receptors

After analyzing the entire human surfaceome and its propensity to be involved in protein-protein interactions, we sought to generate *de novo* protein designs against three major cell surface receptors: fibroblast growth factor receptor 2 (FGFR2), human epidermal growth factor receptor 3 (HER3) and interferon alpha/beta receptor 2 (IFNAR2). MaSIF-site identified one site with a high interface propensity for each targeted structure (**Figure 3A**). All putative interfaces were situated at or near previously identified binding sites (**Supplementary Figure S3**). Using the MaSIF-seed-search pipeline [10], we computationally generated and selected 2,000 protein designs against each target. Briefly, top-ranking seeds were refined with the Rosetta modeling suite [14] and grafted on recipient scaffold protein, generating about 2,000 designs for each target (rd1) and sampling a wide range of folds and topologies (**Figure 3B**).

**Figure 3:**
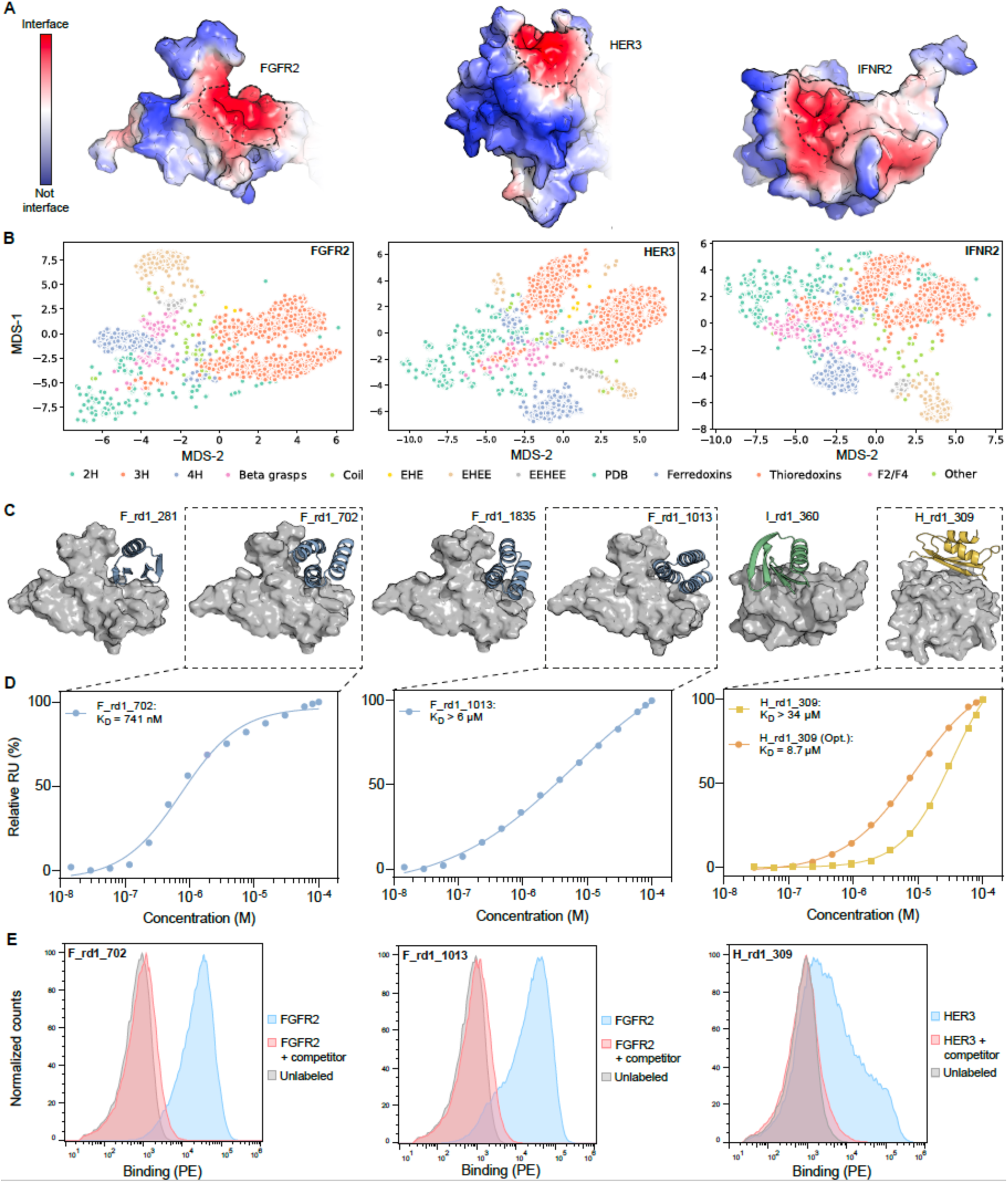
Generation of *de novo* designed binders against surface receptors. **A.** Interface prediction done by MaSIF-site on the three protein targets. Red color indicates a high interface propensity, while blue color indicates a low propensity. **B.** Fold diversity of the ∼2,000 computational designs for the three target proteins mapped using multidimensional scaling (MDS) of pairwise RMSDs between all designs. **C.** Computational models of the 6 experimentally validated designs. Target proteins are shown in gray, and designs are colored in blue (FGFR2 binder), green (IFNAR2 binder) or yellow (HER3 binder). **D.** Affinity measurements of three expressing binders performed by SPR. Steady state responses and dissociation constants were plotted using a nonlinear four-parameter curve fitting analysis. **E.** Histograms of the binding signal (PE, phycoerythrin) measured by flow cytometry on yeast displaying designed binders. Yeast were labeled with their respective target protein in free form (blue) or pre-blocked with a competitor protein (red, 7NIJ for FGFR2 and RG7116 for HER3) or unlabeled (gray).

All computational designs were screened by yeast surface display, and after two rounds of sorting, we observed clear binding populations to their respective targets, which were selected for deep sequencing. We identified 10 designs for FGFR2, 3 designs for HER3, and 1 design for IFNAR2 for which deep sequencing showed an enriched binding profile, confirmed using flow cytometry (**Supplementary Table S1**). Half of the designs showed the predicted binding mode as assessed by mutagenesis at the designed interface (4 for FGFR2, 1 for IFNAR2, and 2 for HER3) (**Supplementary Figure S3** and **Supplementary Table S1**).

Upon expression in *E. coli* (**Figure 3C**) and subsequent purification, these candidate binders were found to adopt dimeric or tetrameric oligomeric states in solution. Evaluation using surface plasmon resonance (SPR) revealed a binding affinity ranging from high nanomolar to low micromolar, as well as a high thermal stability (**Figure 3D**, **Supplementary Figure S4**). Importantly, competition assays showed the absence of binding in the presence of a known competitor, confirming that the correct target interfaces are engaged by the different designs (**Figure 3E**, **Supplementary Figure S3** and **Supplementary Table S1**).

Of note, one of our designs, H_rd1_309, only expressed in presence of a soluble fusion protein (Maltose Binding Protein, MBP) and demonstrated a binding affinity higher than double-digit μM, most probably because of improper folding (**Supplementary Figure S5**). We therefore sought to improve its stability and expressibility by using ProteinMPNN [15], with fixed binder interface residues and filtering the designed sequences with AF2 prediction metrics [11] to retain one candidate (**Supplementary Figure S5**). The prediction of the optimized design done by AF2 multimer [16] also showed perfect alignment with the original model (1.64 Å Cα-RMSD). Finally, the optimized H_rd1_309 expressed in high yield without soluble fusion protein and showed increased affinity on SPR (K_D_ = 8.7 μM). This result, together with similar findings from other groups [8], [9], [17], underscores the need for an optimized pipeline that leverages state-of-the-art deep learning tools to improve and select designed protein binders.

### Enhancing experimental successes with ML tools

Following the observed limited experimental success in terms of binder design success rate (for instance 4/2,000 binders for FGFR) and the binders’ challenging biophysical properties (*e.g.* low expression and stabilities), we set out to improve the MaSIF-seed-search design pipeline by leveraging the machine learning toolbox including ProteinMPNN [14] and AF2 [11]. The previously seen shortcomings could have resulted from the designs failing to: a) fold properly and/or b) bind the targeted site effectively. Within the MaSIF-seed-search pipeline the bulk of the mutations were performed to design the protein interaction site, while the remainder of the scaffold was largely untouched.

Hence, we sought to optimize the MaSIF-seed-search pipeline using ProteinMPNN and assess their in-silico foldability and interface quality using AF2 (**Figure 4A,B**, **Supplementary Table S2**), as a strategy to enhance their biochemical behavior. In the first stage of optimization, ProteinMPNN was used to optimize the designed sequences in two settings: *i*) non-interface residues and *ii*) full redesign of the binders, in an attempt to improve their affinity. The *in silico* foldability of the ProteinMPNN-generated sequences was determined employing AF2 predictions. The designs were filtered based on pLDDT and C_⍺_ RMSD to the original model. The filtered designs’ interface quality was then assessed employing complex prediction using the AF2 monomer model and selecting for candidates with interface pTM (ipTM) ≧ 0.7 [16], [18]. Finally, we tested ∼180 sequences for each of FGFR2 and IFNAR2, and ∼500 sequences for HER3.

**Figure 4:**
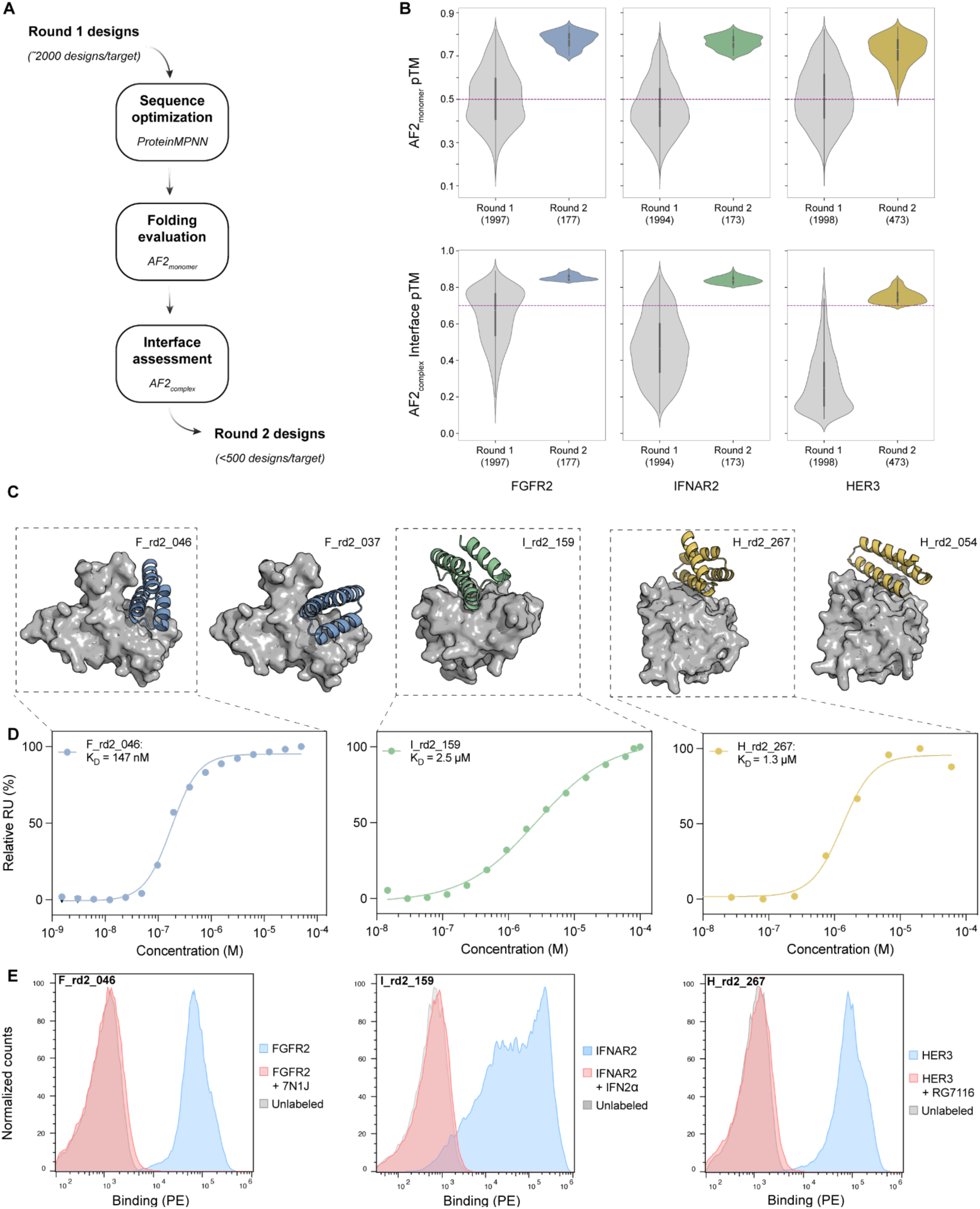
Improved design pipeline and experimental characterization of optimized designs. **A.** Novel computational pipeline including ProteinMPNN for interface design and/or scaffold optimization and AF2 for monomer folding and complex prediction **B.** Comparison between designs before and after optimization for AF2 monomer pTM (top, dashed line set at pTM=0.5) and AF2 multimer ipTM (bottom, dashed line set at ipTM=0.7). **C.** Computational models of 5 experimentally validated optimized designs. Target proteins are shown in gray and protein designs are colored in blue (FGFR2 binder), green (IFNAR2 binder) and yellow (HER3 binder). **D.** Affinity measurements of three binders performed by SPR. Steady state responses and dissociation constants were plotted using a nonlinear four-parameter curve fitting analysis. **E.** Histograms of the binding signal (PE, phycoerythrin) measured by flow cytometry on yeast displaying designed binders testing the binding specificity with competing reagents. Yeast were labeled with their respective target protein in free form (blue) or pre-blocked with a competitor protein (red) or unlabeled (gray).

The optimized designs (rd2) were screened using yeast display, and after two rounds of sorting, yeast cells exhibiting binding signals to their target proteins were selected for deep sequencing. A subset of designs, selected based on enrichments derived from deep sequencing data, was expressed in *E. coli*. All designs were readily expressed except for F_rd2_215 (**Supplementary Table S1**). The purified designs were folded, monomeric, and showed high nanomolar to low micromolar affinities against their respective targets, as determined by SPR, namely F_rd2_046 (K_D_ = 147 nM), I_rd2_159 (K_D_ > 2.5 µM), and H_rd2_267 (K_D_ = 1.3 µM) for FGFR2, IFNAR2 and HER3 respectively (**Figure 4C-D**, **Supplementary Figure S7, S8**). Interface knockout (KO) mutants and competition assays with known binding reagents were consistent with the designed modes of interaction (**Figure 4E**). Our results highlight the key role of the optimized pipeline utilizing ProteinMPNN and AF2 in improving the experimental success rates for identifying site-specific binders, with enhancements exceeding 11-fold and 8-fold compared to rd1 designs for FGFR2 and HER3, respectively. Moreover, while the lead rd1 designs targeting IFNAR2 failed at the expression stage, the optimization process successfully produced a low micromolar binder (**Supplementary Table S4**).

To assess the contribution of ML tools to the experimental successes of the rd2 designs, we compared the *in silico* metrics of the designs before and after optimization. The rd2 design set showed a much higher *in silico* folding propensity as evaluated by AF2 as compared to the rd1 designs (**Figure 4B** top, **Supplementary Figure S9A**, **Supplementary Table S3**). The key discriminating metric was the interface quality, inferred from the ipTM score, where most of the rd1 designs were below 0.7 (**Figure 4B** bottom, **Supplementary Table S3**). These findings align with our hypotheses that the lower success rates of rd1 designs were likely due to suboptimal protein scaffolds hindering folding and consequently binding. Sequence similarity analysis revealed that more than half of the sequence were altered in most rd2 designs compared to their predecessors (**Supplementary Figure S9D**).

Of the six lead candidates, two were obtained through the optimization of the scaffold while retaining the original interface, and four resulted from the fully redesigned sequences. Although the rd2 AF2 complex predictions closely aligned with the rd1 models, showing only minor structural deviations on the binders (**Supplementary Figure S10A,B**), the rd2 designs demonstrated improved AF2 metrics for both *in silico* folding and complex interface quality compared to their respective rd1 designs (**Supplementary Figure S10C**). These results highlight the capabilities of AF2 to aid in protein design tasks.

### Generation of peptide binders from *de novo* proteins

Building on the success of mini-protein binders targeting FGFR2, IFNAR2, and HER3, we expanded our approach to design peptide binders, which offer additional advantages as modality due to the smaller size. Two design strategies were utilized: (i) cyclized peptides generated by diffusion-based methods and (ii) stabilized PPI motif peptides derived from the lead candidates of the mini-protein binders using rational protein design techniques (**Figure 5A**).

**Figure 5:**
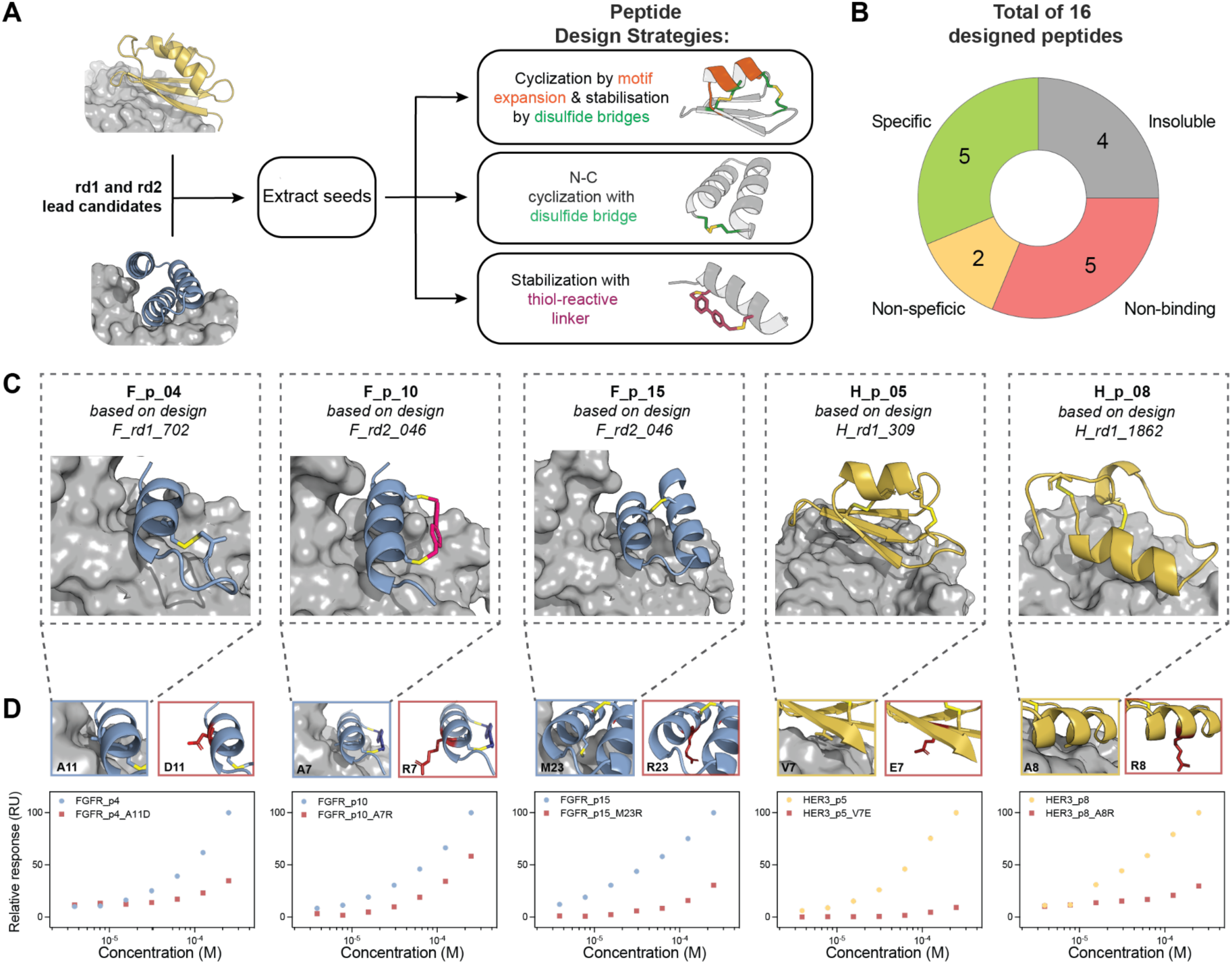
Experimental characterization of stabilized peptides derived from de novo designed mini binders. **A.** Overview of the employed peptide design strategies. The interface binding motifs from the experimentally validated rd1 and rd2 binders were extracted and used to design cyclized peptides. Three types of designs were generated, cyclic peptides with disulfides, peptides with a single disulfide, and peptides stabilized with thiol-reactive linkers. B. Overview of properties of tested peptides. From a total of 16 synthesized peptides, 4 were insoluble, 5 did not show any binding to their target protein by SPR, 4 showed non-target specific binding, and 5 were target- and mode-specific binders. **C.** Computational models of the five experimentally validated peptides targeting FGFR2 (blue) HER3 (yellow). **D.** Affinity and specificity measurements of the five lead peptide candidates were performed using SPR. Maximum response at given concentrations is shown, highlighting specific peptide-target binding (FGFR2:blue, HER3:yellow) and their corresponding interface knockout mutant (red), which exhibit reduced binding signal, thus demonstrating the peptides’ target specificity.

Starting from the experimentally validated rd1 and rd2 designs, the binding interface motifs were analyzed for their suitability to design cyclized peptides. Defined secondary structural motifs such as helix, helix-turn-helix and beta-hairpins were amenable to stabilization through disulfide bonds, achieved either by introducing terminal cysteine residues or incorporating disulfide-anchored loops. In some instances, disulfide stabilized motifs were further embedded into larger cyclized peptides (up to 38 residues in length) featuring two disulfide bonds and employing the cyclotide design pipeline. The cyclized peptides were evaluated *in silico* for their predicted folding, using AF2 with relative positional encoding adjusted to account for inter-residue distances in a cyclic sequence [19], and for the stability of the engineered disulfide bonds, using molecular dynamics simulations. Additionally, a minimalistic approach was employed, wherein helical interface motifs were extracted and stabilized through the addition of a thiol-reactive linker (**Figure 5A**). A total of 16 peptide candidates were designed, synthesized, and experimentally tested, resulting in 5 target-specific binders (**Figure 5B**).

12 of the 16 peptides were soluble in PBS, with four showing folded structures in solution as assessed by CD spectroscopy: two peptides (F_p10 and F_p15) in PBS and two (F_p1 and H_p8) in the presence of 20% TFE (2,2,2-Trifluoroethanol) (**Supplementary Figure S11, Supplementary Table S5**). Binding of the 12 peptides was tested with SPR against their respective targets (**Figure 5C**). Target specificity for the designed peptides was determined by SPR against control proteins and by interface knockout mutants against their targets (**Figure 5C**, **Supplementary Figure S12**). We identified five target-specific peptides, three targeting FGFR2 (F_p4, F_p10, and F_p15) and two targeting HER3 (H_p5 and H_p8) (**Figure 5C, Supplementary Figure S12)**.

## Discussion

In this study, we employed a surface-based geometric deep learning approach (MaSIF) to systematically predict and analyze protein binding sites across the entire human surfaceome. This extensive analysis unveiled approximately 4,500 sites that are potentially targetable by protein-based ligands, opening new routes to modulate the function of these cell-surface receptors. To follow up on the prediction of binding sites across the surfaceome, we utilized a large library of 640,000 protein seeds to provide *bona fide* motifs to engage the targets of interest. This effort, equivalent to conducting 3 billion protein-protein docking and scoring simulations, provided detailed insights into the targetability of each predicted binding interface and identified a high-quality set of binder seeds to support future target-specific protein design initiatives. This depth of structural exploration is out of reach to other design methods relying on diffusion [9] or structure prediction networks [20]. To experimentally validate the practical utility and applicability of the selected seeds, we designed binders to target three proteins using two modalities — mini-binders and stabilized peptides, both of which demonstrated a high success rate when initiated with the seeds identified through surface fingerprinting with MaSIF.

Several recent efforts have focused on *de novo* protein binder design [9], [10], [20], [21]. Having a defined target site on a protein of interest, both physics-based [5] and deep learning methods such as RFdiffusion [9] or BindCraft [20] can provide unconditional designs with various success rates. Despite this progress, challenges remain in searching the vast space of possible binding modes and accurately identifying binding sites with high potential for targetability. Many current approaches lack for instance prior information about binding site properties or structural templates, leading to suboptimal starting points and potentially low experimental success rates. One way to address this issue is by leveraging high-quality preliminary data, including detailed structural information about binding sites and curated initial binder seeds which MaSIF can leverage at a proteome-wide scale. As demonstrated in the current study, such data can serve as a foundation to reduce the design space, improve prediction accuracy, and increase the likelihood of successful experimental outcomes. In this work we also explored how to best use machine learning tools to enhance binding activity of designed binders, contributing to the general advancement of the protein design field for the engineering of molecular recognition. It is, however, important to mention that it remains challenging to computationally design high affinity binders.

Furthermore, we introduced SURFACE-Bind (https://surface-bind.inria.fr), a first-of-its-kind large-scale interactive webserver, which allows the user to explore the analyzed target proteins and their top-scoring seeds for each predicted binding site, making it a valuable resource for biologically-driven questions as well as for drug discovery efforts. This platform facilitates effortless browsing and data retrieval for specific target proteins. While this study focused on the human surfaceome, the database can be expanded to include surface proteins from other organisms, fostering advancements in protein binder design across diverse research efforts, including those with therapeutic applications.

## Data availability

All data, including comprehensive details of binding sites, unbound-state scores, predicted seeds, and bound-state scores, are freely accessible on Zenodo (link to be added). Additionally, the SURFACE-Bind web server (https://surface-bind.inria.fr/) offers an interactive platform for exploring predicted binding sites and their corresponding seeds.

## Code availability

The code and scripts for reproducing the results of this study are freely available online. MaSIF-site can be accessed at https://github.com/LPDI-EPFL/masif, while the seed search protocol, based on MaSIF-seed-search, is available at https://github.com/LPDI-EPFL/masif_seed. Additionally, all scripts for executing the workflow on a large scale, defining unbound-state scores for binding sites, analyzing the results, and optimizing binders resulting from MaSIF-seed-search can be found at https://github.com/hamedkhakzad/SURFACE-Bind.

## Acknowledgment

We thank SCITAS at École Polytechnique Fédérale de Lausanne (EPFL) for support in running our pipeline and all design trajectories; Bertrand Wallrich and Frederic Beck (Service d’expérimentation et de développement (SED) at Centre Inria de l’Université de Lorraine) for support in running SURFACE-Bind web server. HK was supported by the French Agence Nationale de la Recherche (ANR), under grant ANR-22-CPJ2-0075-01. AM was supported by the Swiss National Science Foundation with grant 310030_197724 and the National Center of Competence in Research in Molecular Systems Engineering with grant 182895. BEC was supported by the Swiss National Science Foundation through a ERC Consolidator replacement grant. The project was also supported by the NovoSTAR program of Novo Nordisk.

## Contributions

S.F., H.K., C.Y. and B.E.C. conceived the idea and supervised the project. H.K. developed and performed the large-scale analysis of the human surface proteome and the seed-searching. P.E.M.B, A.M. and A.S conducted experimental characterization. T.Y.Y, and J.D. developed the workflow for cyclotide design and performed the computational peptide design. A.S. and A.M. performed the computational design of *de novo* mini-protein binders. A.S. designed the cyclized and thiol-reactive linker peptides. H.K. developed the web server for the SURFACE-bind and organized the GitHub repository. All authors contributed to the analysis of the results. H.K., P.E.M.B., A.S., A.M., T.Y.Y., J.D. and B.E.C. wrote the paper with input from all co-authors.

## Ethics declarations

## Methods

### Computational workflow

The surfaceome database was collected from SURFY [2], where 2,886 proteins were predicted *in silico* to be on the surface of human cells. Corresponding AF2 models for each entry were obtained and filtered with the criteria of pLDDT>0.6. MaSIF-site [12] was then used to predict the binding sites for all proteins and the results were clustered using DBSCAN, employing 0.7 as threshold for a point to be considered as a binding site. As each atom was sampled by 10 points, we discarded binding sites with a number of points < 200 (1-3 residues coverage) and those located in transmembrane domains, resulting in a total number of 4,500 binding sites. The following geometric and chemical features of each predicted binding site were computed: shape index, curvature (concave, convex, flat), area in A^2^ (geometrical), sequence/chemical composition, solvent accessible surface area (SASA), and the hydrophobicity scale of Kyte and Doolittle (chemical). The bound state scores were calculated per seed through the “binding seed identification” workflow. All scripts and notebooks to run the workflow are available at: https://github.com/hamedkhakzad/SURFACE-Bind.

### Binding seed identification

The fingerprint of the 12 Å (geodesic radius) patches found at the predicted interface of each target site was used to find complementary fingerprints in the MaSIF-seed database. The database consists of ∼640,000 continuous structural fragments (seeds), amounting to 402 million surface patches/fingerprints [10]. The MaSIF-search algorithm [12] was trained to make patch fingerprints similar for interacting patches and dissimilar for non-interacting patches. As described previously [10], seeds are selected based on a high interface propensity score and a low fingerprint distance (Euclidean distance between target and seed fingerprint). In a second-stage alignment and scoring using the RANSAC algorithm, seeds were selected based on the interface post-alignment (IPA) score. The binding seed identification was applied to all 4,500 sites, performing nearly 3 billion pairwise docking runs to obtain the final list of seeds per target site.

### Seed and interface refinement

As done previously [10] to optimize the binding energy of seeds *in silico*, a FastDesign protocol in Rosetta [22] was employed, incorporating a penalty for buried unsatisfied polar atoms in the scoring function [23]. Optimized seeds were selected based on several criteria: computed binding energy (ddG), shape complementarity, the number of interface hydrogen bonds, the number of buried unsatisfied polar atoms, and the number of atoms in contact with the small molecule. Beta sheet-based motifs making more than 33% contact with the target complex using loop regions were discarded. Additionally, the uniqueness of each seed was assessed through pairwise alignment of hotspot residues. For seeds showing more than 70% hotspot identity with another seed, only the one with the best surface-normalized ddG was retained.

### Seed grafting

Selected seeds were then grafted using the Rosetta MotifGraft protocol [24] to stabilize the binding motif and enhance contact with the target complex as done in [10]. Each seed was matched with a database of approximately 6,500 small protein scaffolds (under 90 amino acids) derived from small globular monomeric proteins in the Protein Data Bank (PDB) [25] and four miniprotein databases that had been experimentally validated [26], [27], [28], [29].

Seeds were trimmed to the minimum number of residues contacting the target, and loop motifs were removed from beta sheet-based seeds to optimize grafting success. After grafting, scaffolds underwent sequence optimization using a similar FastDesign protocol in Rosetta as for the seed refinement. Final designs were selected based on computed binding energy (ddG), shape complementarity, the number of interface hydrogen bonds, and the count of buried unsatisfied polar atoms.

### Structural and sequence similarity calculations

To calculate the structural similarity between the AF2 prediction and the initial designs, the C_⍺_ RMSD was calculated as the mean Euclidean distance between the aligned structures via Biopython package [30]. The sequence similarity between two given sequences was calculated using the Biotite python package (v. 0.41) [31], [32].

### Sequence optimization using ProteinMPNN

We used a ProteinMPNN model with soluble MPNN (MPNN_sol_) weights, trained on a dataset comprising only soluble proteins [33]. The designs generated in round 1 were utilized as inputs to ProteinMPNN. Sequence optimization was performed in two settings: (i) **Fixed interface**, where interacting residues of the designs within 3.5 Å from the target interface were kept fixed while ProteinMPNN redesigns the scaffold residues, and (ii) **Full redesign**, where all residues were optimized. For each design, 50 sequences were generated, 25 sequences with each setting. From each setting, the top 5 unique sequences per design, based on ProteinMPNN global score, were forwarded to AF2 for subsequent filtering steps.

### Design filtering using AlphaFold2

The *in silico* foldability of the ProteinMPNN-generated sequences was assessed using AlphaFold2 via ColabFold (v1.5.2) [34]. Predictions were performed with three recycles and in single sequence mode (no multiple sequence alignments were provided). The AF2 scores from the 5 prediction models were averaged and used to filter the optimized sequences based on the following criteria: (1) pLDDT (predicted local distance difference test) ≧ 90 except for HER3, where the threshold was lowered to ≧ 80, and (2) C_⍺_ RMSD to design model ≦ 1.5 Å.

The binder-target complex interface of the passing designs was further evaluated using complex prediction performed with AlphaFold2 monomer model via ColabDesign (v1.1.1) [37]. Custom templates of the target-binder complex were provided for these predictions. The designs were subsequently filtered based on their ipTM (interface predicted TM-score) ≧ 0.7. Finally, a selection of successful designs was chosen for experimental testing.

### Library preparation

For each target complex - namely FGFR2, IFNAR2 and HER3 - around 2,000 designs were reverse-translated into DNA and purchased from Twist Bioscience as oligo pools with 18 base pairs homology overhangs with the final vector. These oligo pools were subjected to two rounds of PCR: first, to amplify the library using the 18 base pairs overhangs, and second, to add 45 base pairs homology with the yeast display vector (with an annealing temperature of 57.5°C for 30 seconds, and an extension at 72°C for 1 minute, over 15 cycles). The amplified inserts and linearized pCTcon2 vector (with C-terminal HA-tagged) were used to transform EBY-100 yeast by electroporation, following previously established protocols [35]. The transformed yeast cells were cultured in minimal glucose (SDCAA) medium at 30°C and then induced overnight in minimal galactose (SGCAA) medium before sorting.

### Flow cytometry analysis and sorting

Induced yeast cells were rinsed with PBS containing 0.1% BSA and then labeled with the target binding molecule (with Fc-fusion) for 2 hours at 4°C. The cells were subsequently pelleted and labeled with a FITC-conjugated goat anti-HA tag antibody (Bethyl; ref: A190-138F; display tag; 1:100 dilution) and a PE-conjugated goat anti-human Fc antibody (Invitrogen; ref: 12-4317-87; binding tag; 1:100 dilution) for 30 minutes at 4°C. Finally, pelleted cells were resuspended in an appropriate PBS/BSA buffer volume and analyzed using a Gallios flow cytometer (Beckman Coulter) or sorted with a Sony SH800 cell sorter. Data acquisition was performed using the Kaluza software (Beckman Coulter, v.1.1.20388.18228) for flow cytometry and the LE-SH800SZFCPL Cell Sorter software (Sony, v.2.1.5) for fluorescence-activating cell sorting. Each designed library was sorted into a binding and a non-binding population. The flow cytometry data were then analyzed with the FlowJo software (BD Biosciences, v.10.8.1).

### Library sequencing

Sorted yeast were cultured, and plasmids encoding protein designs were extracted using the Zymoprep Yeast Plasmid Miniprep II kit (Zymo Research) according to the manufacturer’s instructions. The gene of interest was then amplified by PCR using vector-specific primers flanking the protein design gene. A second PCR was conducted to add Illumina adapters and Nextera barcodes. The PCR product was then desalted and purified using the Qiaquick PCR purification kit (Qiagen). Next-generation sequencing was performed using the Illumina MiSeq system with 500 cycles. Approximately 0.8-1.2 million reads per sample were obtained, translated into the appropriate reading frame, and matched with the expected input sequences from the libraries. The enrichment of each design was calculated by dividing the counts in the binding population with those in the non-binding populations. Finally, hits were identified if the enrichment in the binding population was more than 10-fold greater than in the non-binding population, and the number of counts in the binding population exceeded 10,000.

### Yeast surface display of single designs

Genes encoding single designs were obtained from Twist Bioscience, featuring ∼25 base pairs homology overhangs for cloning. Each design was inserted into pCTcon2 plasmids (with C-terminal HA-tagged) via Gibson assembly and transformed into HB101 competent bacteria for DNA production. After purification and sequence verification by Sanger sequencing, the DNA was used to transform competent EBY-100 yeast with the Frozen-EZ Yeast Transformation II Kit (Zymo Research). Transformed yeast cells were cultured in minimal glucose (SDCAA) medium at 30°C and then induced overnight in minimal galactose (SGCAA) medium before undergoing flow cytometry analysis.

### Protein expression and purification

Protein sequences used in this work are listed in **Supplementary Table S6.** Genes encoding the 6xHis-tagged and/or human Fc-tagged proteins were obtained from Twist Bioscience, cloned into pET11 (bacterial vector) or pHLSec (mammalian vector) using Gibson assembly, and transformed into HB101 bacteria. Plasmids were extracted using a GeneJET plasmid Miniprep kit (ThermoFisher) for bacterial vectors or a Plasmid Plus Midi Kit (Qiagen) for mammalian vectors and verified by Sanger sequencing. Proteins were then purified using either bacterial or mammalian expression systems.

For mammalian expression, the Expi293 system (ThermoFisher; ref: A14635) was used. Supernatants were collected after 6 days, filtered, and purified as described below. For bacterial expression, BL21(DE3) or T7 Express Competent *E. coli* were transformed with the plasmid and grown overnight in a pre-culture. Pre-cultures were then inoculated 1:50 in 500 ml medium and incubated at 37°C until reaching an OD600 of ∼0.7. The cultures were induced with 1mM IPTG and incubated overnight at 18-20°C. Cells were harvested by centrifugation at 4000g for 10 minutes, resuspended in lysis buffer (50 mM Tris, pH 7.5, 500 mM NaCl, 5% glycerol, 1 mg/ml lysozyme, 1 mM PMSF, and 1 µg/ml DNase), and lysed by sonication. Lysates were clarified by centrifugation at 30,000g for 30 minutes and filtered. All 6xHis-tagged proteins were purified using the ÄKTA pure system (GE Healthcare) with a Ni-NTA HisTrap affinity column, followed by size exclusion chromatography on a Superdex HiLoad 16/600 75pg or 200pg column, depending on the protein size. Proteins were concentrated in PBS as the final buffer.

### Surface plasmon resonance

Affinity measurements were conducted using a Biacore 8K system (GE Healthcare, software v4.0.8.19879) with HBS-EP+ as the running buffer (10 mM HEPES at pH 7.4, 150 mM NaCl, 3 mM EDTA, 0.005% v/v Surfactant P20, GE Healthcare). Proteins were immobilized on a CM5 chip (GE Healthcare #29104988) via amine coupling to achieve 500–1500 response units (RU). Analytes were injected in serial dilutions using the running buffer, with a flow rate of 30 μL/min for a contact time of 120 seconds, followed by a dissociation time of 400 seconds. The SPR data were analyzed by steady-state affinity mode, reporting the RU normalized to the highest RU reached for each concentration.

### Size-exclusion chromatography–multi-angle light scattering (SEC-MALS)

To determine the molecular weight of the purified designs, SEC-MALS on a miniDAWN TREOS instrument (Wyatt) was utilized. The samples, at a concentration of approximately 1 mg/ml in PBS (pH 7.4), were injected into a Superdex 75 10/300 GL column (GE Healthcare) with a flow rate of 0.5 ml/min. Signals for UV_280_, refractive index (dRI), and light scattering were recorded. The molecular weight was calculated using the ASTRA software (version 6.1, Wyatt).

### Circular dichroism

Far-ultraviolet circular dichroism spectra were recorded using a Chirascan spectrometer (Applied Photophysics). Protein samples were diluted in PBS to a concentration of 300 μg/ml and placed in a 1 mm path-length cuvette. Wavelengths between 200 nm and 250 nm were scanned at a speed of 20 nm/min with a response time of 0.125 s, with all spectra corrected for buffer absorption. Temperature ramping melts were conducted from 20 to 90 °C at a rate of 2 °C/min. Thermal denaturation curves were plotted by monitoring changes in ellipticity at the global minimum, and the melting temperature (T_m_) was determined by fitting the data to a sigmoid curve equation using GraphPad Prism.

### Stapling of peptides with a chemical linker

The peptides were chemically synthesized using solid phase peptide synthesis and the chemical linker was introduced following the procedure described by Diderich et. al. [36] with some modifications. The thiol-reactive linker employed in this study is 4,4’-Bis(bromomethyl)biphenyl. The synthesized peptide was incubated with the chemical linker in the presence of a reducing agent (TCEP) with 1:1.5:3 molar ratios respectively. The reaction was incubated for 120 minutes at 25°C and 1000 rpm in a thermomixer. Finally, the stapled peptide was purified using RP-HPLC and lyophilized until further use. This procedure was performed by GL biochem (Shanghai), Ltd., where the peptides were purchased from.

### Generation of the disulfide fragment hashtable

N-to-C terminus disulfide monocycles ranging from 6 - 15 mers were generated using the kinematic closures algorithm in Rosetta [27]. Briefly, starting with polyglycine with N and C terminal cysteines, backbone torsions were biasedly sampled based on rama probability. Solutions with ideal disulfide chi angles are clustered based on backbone torsion bin strings [37]. To generate the hashtable, sliding windows containing at least 3 residues containing both cysteines were taken. The hash key is the affine transform between N-C_α_-C local frames at perspective ends and the value corresponds to the internal degree of freedom of the peptide fragments [38].

### Cyclotide design of disulfide-dense cyclized peptides

Binding motifs were extracted from experimentally validated round 1 (rd1) designs. A disulfide anchored loop was added to the selected motifs through disulfide fragment lookup using the hashtable generated from disulfide monocycles. During the lookup, one external N-C_α_-C and one internal N-C_α_-C frame were sampled to identify potential disulfide placements while extending the motif. Next, the motifs were stabilised through incorporation into larger - up to 38 residues - cyclic peptides employing RFDiffusion [9]. Briefly, an artificial chain break within the linear peptide was defined, and standard RFDiffusion structure generation was used to cyclize the peptide termini. To further enhance the peptide’s stability, a second disulfide bond was introduced via Rosetta [27] when feasible. The cyclized peptides were evaluated using an AlphaFold2-based cyclic peptide and protein structural prediction network [39]. Predictions were filtered based on the following criteria: (1) pLDDT (predicted local distance difference test) > 80, (2) ipAE (interaction predicted alignment error) < 5, and (3) C_⍺_ RMSD to design model < 2.5 Å. Filtered designs were then assessed using molecular dynamics simulations for their fold stability.

### Molecular dynamics (MD) simulation protocol

MD simulations were performed in explicit TIP3P solvent using the GROMACS 2023 package [40] and the RSFF2 force field [41]. RSFF2 is an AMBER-derived force field which was parameterized using a coil library and shown to capture intrinsic conformational preferences of amino acids with high accuracy. Initial structure predictions for 29 representative designs were solvated in a cubic box of pre-equilibrated solvent, with a minimum distance between the peptide and the box boundaries set to 1.0 nm. The solvated structure was energy-minimized using the steepest descent algorithm and then equilibrated in four stages: 50 ps in the *NVT* ensemble at 300 K, 50 ps *NPT* at 300 K and 1 bar, both with position-restrained heavy atoms; 100 ps *NVT* at 300 K, 100 ps *NPT* at 300 K and 1 bar, both unrestrained. An *NPT* production run was conducted for 250 ns, at 300 K and 1 bar, with a simulation time step of 2 fs and data recorded every 1 ps.

All simulations were performed using the standard leapfrog algorithm, with SETTLE used to maintain water geometry [42], and LINCS applied to constrain hydrogen-containing bonds to equilibrium lengths [43]. Non-bonded interactions were treated explicitly up to 1.0 nm, with Coulombic interactions past this threshold computed via particle mesh Ewald summation, using a 0.12 nm Fourier spacing and cubic interpolation. Dispersion corrections for both energy and pressure were applied to the long-range van der Waals interactions. The V-rescale thermostat with a coupling time constant of 0.1 ps was used to control temperature, and the Parrinello-Rahman barostat with a coupling time constant of 2.0 ps and isothermal compressibility of 4.5×10^−5^ bar^−1^ was used to maintain pressure [44].

All systems were simulated with disulfide bonds replaced with free-sulfhydryl cysteines, and the pairwise distances for all possible pairs of cysteine sulfur atoms were recorded for each frame of the simulation. Pseudo-disulfide frames were defined as all frames with any S–S distance lower than 4 Å. For each system, the overall population of such frames, as well as the relative population of all represented combinations of spatially proximate cysteine pairs, was recorded. Systems with a single dominant combination matching the original design were considered “more synthesizable” and proceeded to the next stage of testing, under the assertion that their conformational ensembles would minimize undesirable side products being formed during synthesis. In addition, to ensure that the stereochemistry of the disulfide bonds would match the original design, we also monitored the distribution of the torsion angles corresponding to the pseudo-disulfides throughout the simulation trajectories. Any systems displaying multimodal distributions, or significant peaks not matching the originally predicted structure were discarded at this stage, whereas the remaining systems proceeded to experimental evaluation.

## Supplementary materials

**Supplementary Figure 1:**
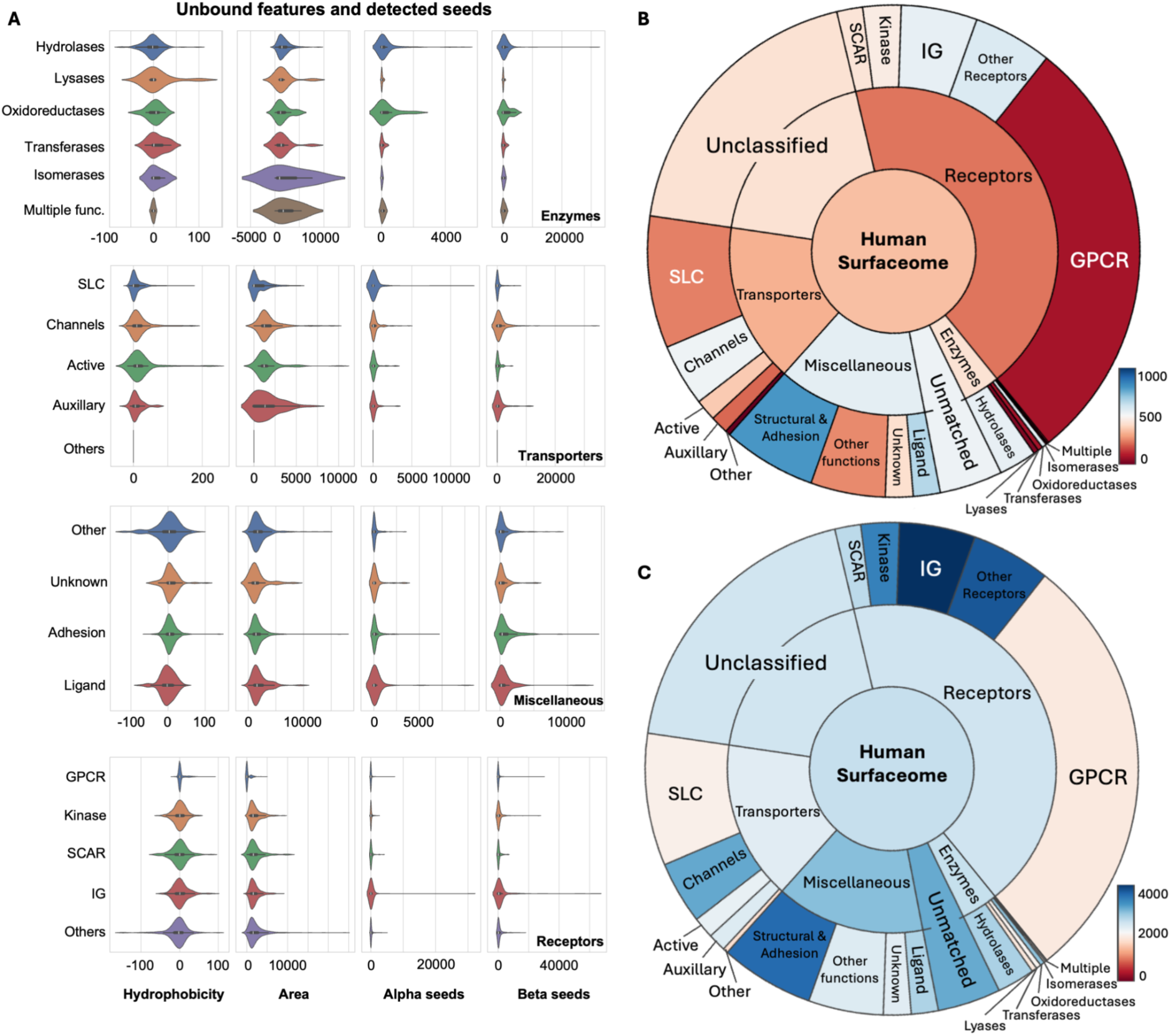
The distribution of unbound features and detected seeds across protein families. **A.** The distribution of two unbound features (hydrophobicity and area of the binding sites) and detected alpha and beta seeds for protein subfamilies (the negative values for “Area” are due to kernel density estimation of the violin plot). **B-C.** The average number of detected alpha and beta seeds across all protein families.

**Supplementary Figure 2:**
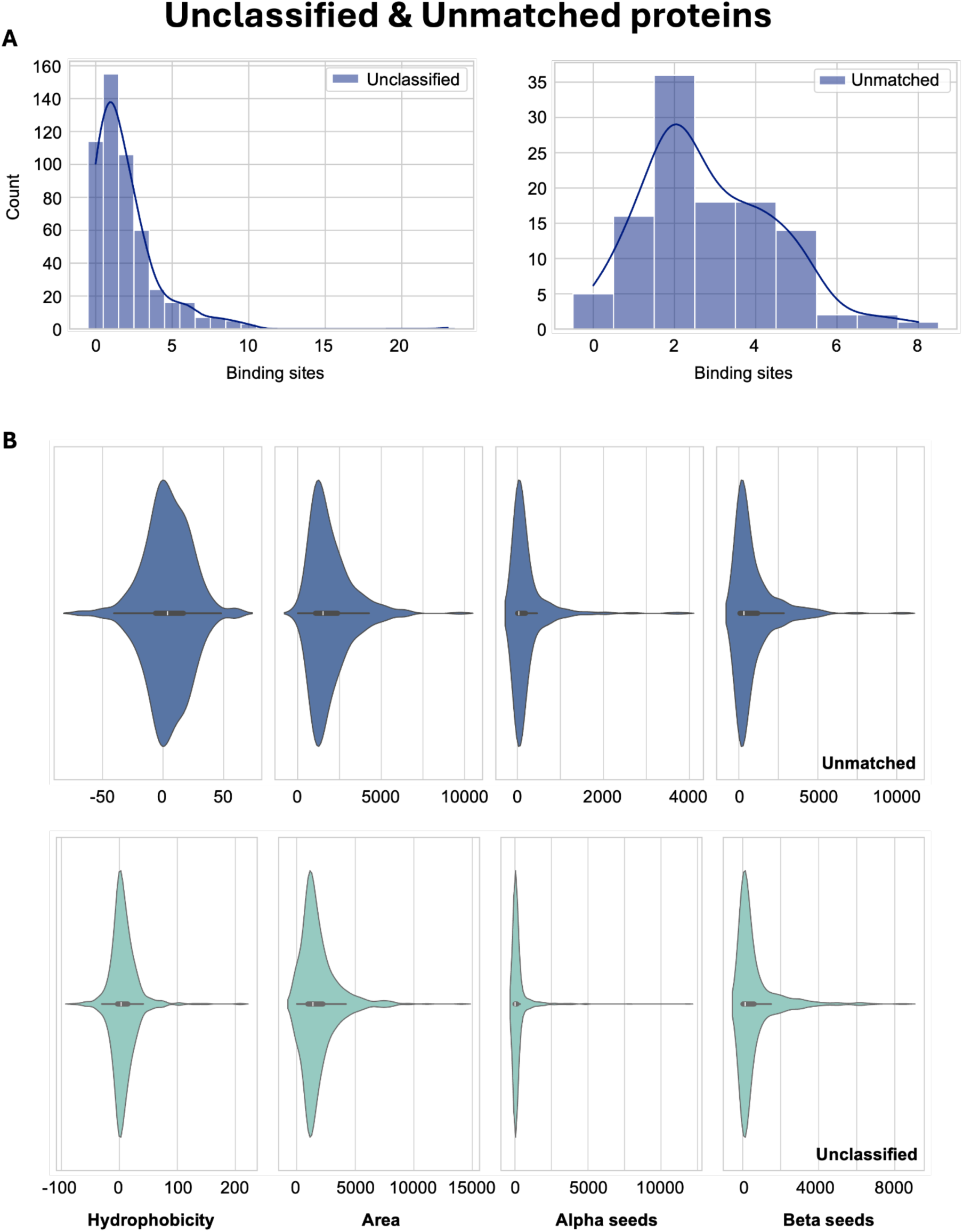
More details for unclassified and unmatched main classes. **A.** The binding site histogram. **B.** The distribution of important unbound features (hydrophobicity and area), and number of detected alpha and beta seeds.

**Supplementary Figure S3:**
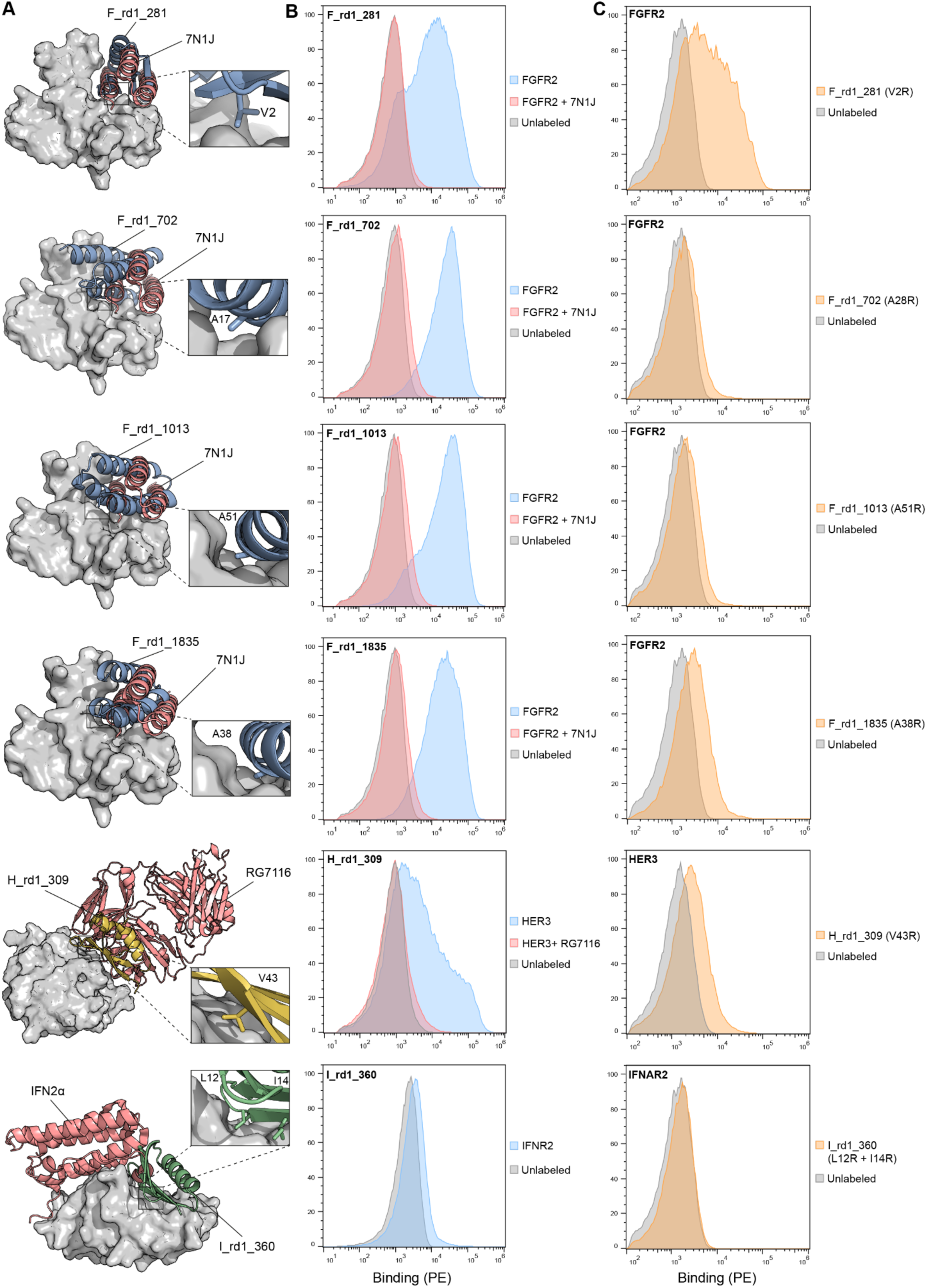
Competition assays and interface point mutation of designed binders (round 1). **A.** Computational model of designed binders (Blue, yellow or green) in complex with their respective target protein (Gray) and overlaid with a competitor (red). Competitors are a computational designed binders (PDB ID: 7N1J), RG Fab (PDB ID: 4LEO) or an Interferon alpha 2 (PDB ID: 2LAG) for FGFR2, HER3 and IFNAR2 respectively. **B.** Histograms of the binding signal (PE, phycoerythrin) measured by flow cytometry on yeast displaying designed binders. Yeasts were labeled with their respective target protein in free form (blue) or pre-blocked with a competitor protein (red) or unlabeled (gray). **C.** Histograms of the binding signal (PE, phycoerythrin) measured by flow cytometry on yeast displaying designed binders with a point mutation at the predicted interface. Yeasts were labeled with their respective target protein (orange) or unlabeled (gray).

**Supplementary Figure S4:**
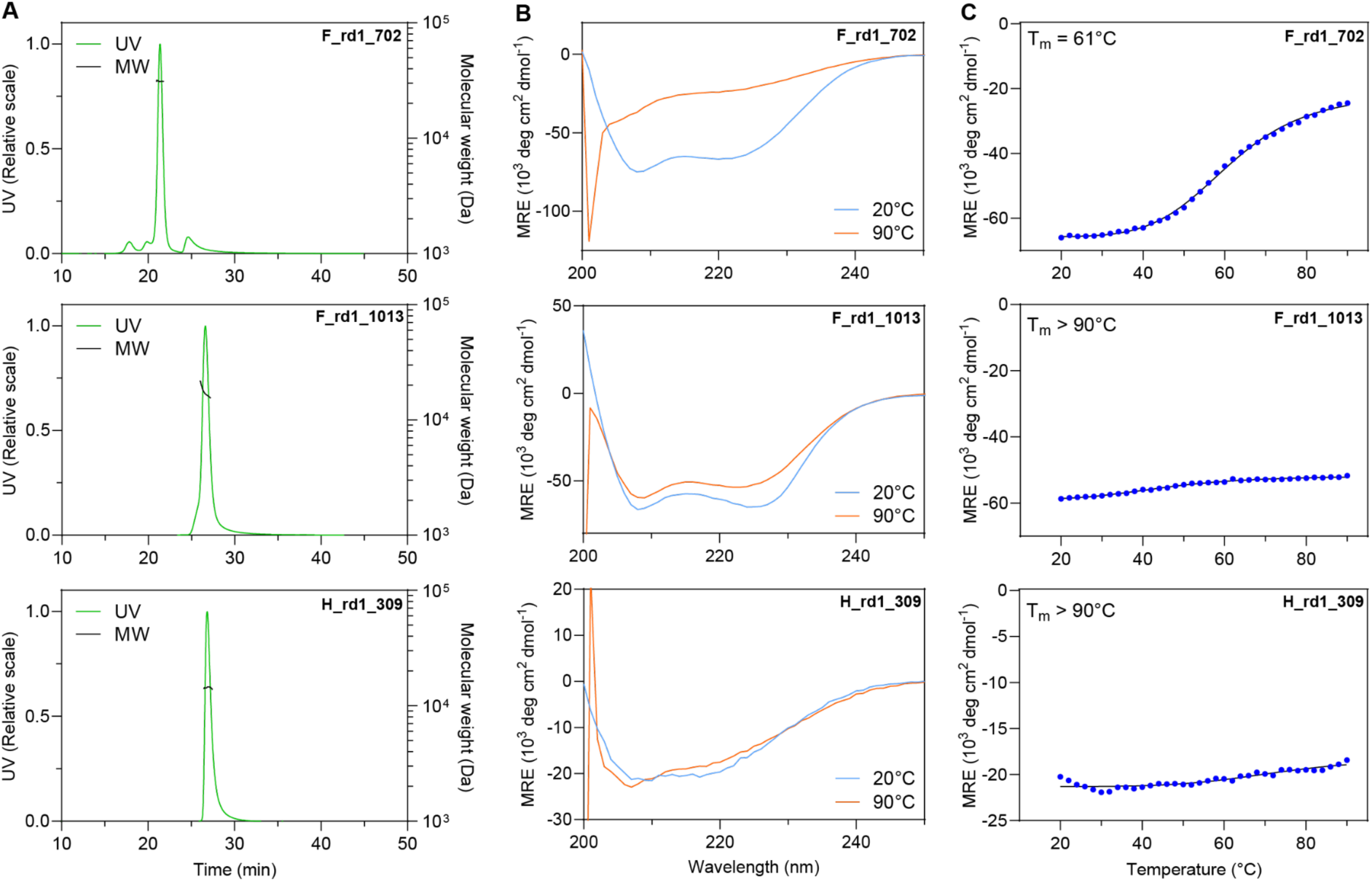
Biophysical characterization of purified binders (round 1). **A.** Oligomeric state determined by size-exclusion multi-angle light scattering (SEC-MALS). **B.** Protein folding of the purified binder measured by circular dichroism at 20°C (blue) or 90°C (orange). **C**. Thermal stability determined by measuring the ellipticity at 218 nm at increasing temperature.

**Supplementary Figure S5:**
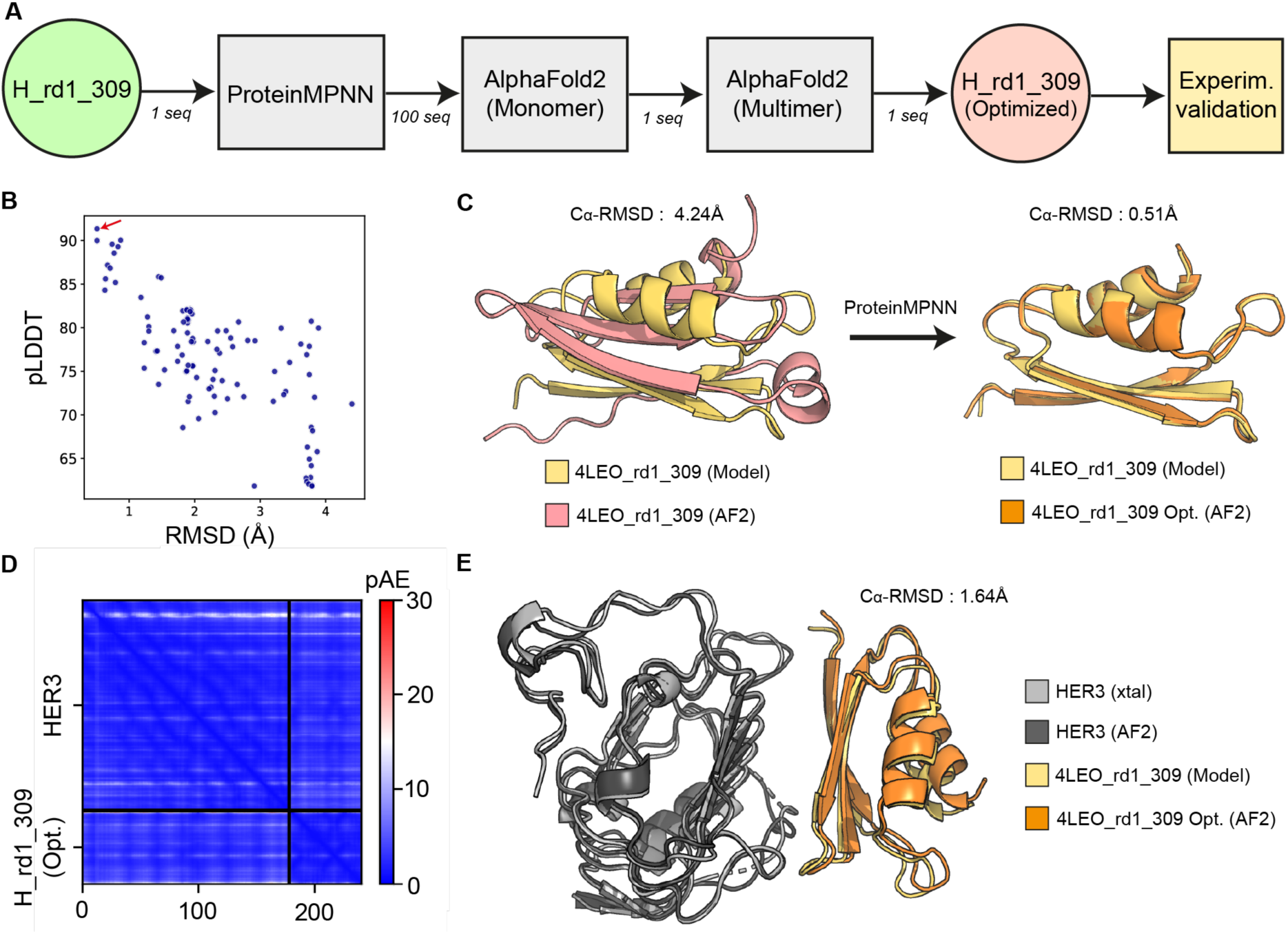
Computational optimization of H_rd1_309. **A.** Computational pipeline for the optimization of the H_rd1_209 binder. The binder sequence was optimized using ProteinMPNN with the binder interface residues kept fixed. The optimized sequences were predicted using AF2 in single-sequence mode and ranked based on overall model confidence, measured by the Predicted Local Distance Difference Test (pLDDT), and Cα root mean-squared deviation (RMSD) relative to the initial model. Subsequently, AF2 multimer was employed to evaluate the complex formation of the top optimized design. Finally, the selected sequence was experimentally validated. **B.** pLDDT and RMSD of the optimized sequences’ AF2 predictions compared to the initial design model. The arrow indicates the design selected for experimental testing. **C.** Structural comparison of the AF2 predictions for the initial design (red) and the optimized design (orange) with the Rosetta model (yellow). **D**. Predicted aligned error (pAE) of the experimentally tested optimized design from the AF2 multimer prediction. **E.** Structural alignment of the AF2 multimer prediction for the selected optimized sequence to the initial Rosetta model (AF2 multimer pTM of 0.9).

**Supplementary Figure S6:**
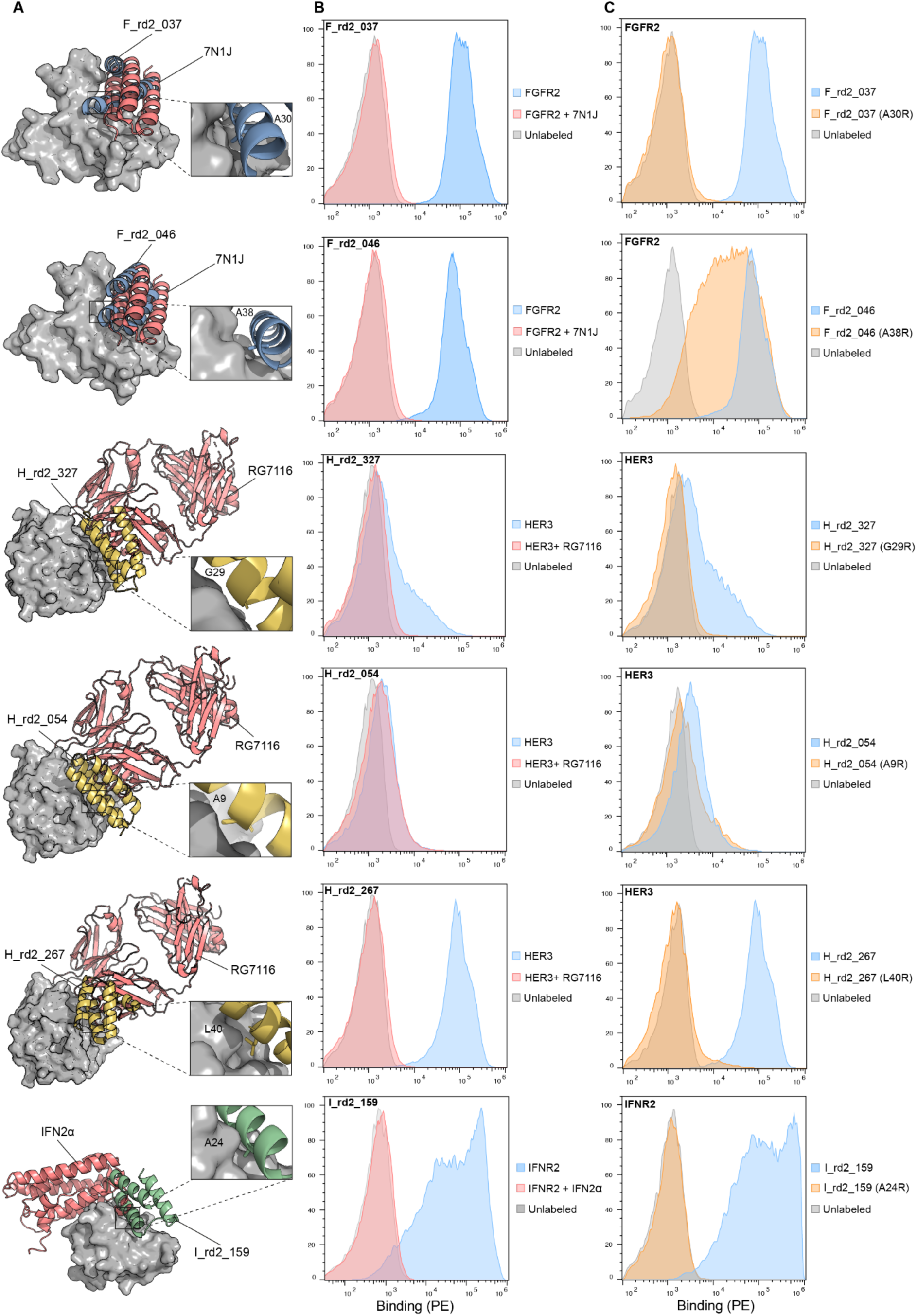
Competition assays and interface point mutation of designed binders (round 2). **A.** Computational model of designed binders (Blue, yellow or green) in complex with their respective target protein (Gray) and overlaid with a competitor (red). Competitors are a computationally designed binder (PDB ID: 7N1J), RG Fab (PDB ID: 4LEO) or interferon alpha 2 (PDB ID: 2LAG) for FGFR2, HER3 and IFNAR2 respectively. **B.** Histograms of the binding signal (PE, phycoerythrin) measured by flow cytometry on yeast displaying designed binders. Yeasts were labeled with their respective target protein in free form (blue) or pre-blocked with a competitor protein (red) or unlabeled (gray). **C.** Histograms of the binding signal (PE, phycoerythrin) measured by flow cytometry on yeast displaying designed binders with a point mutation at the predicted interface. Yeasts were labeled with their respective target protein (orange) or unlabeled (gray).

**Supplementary Figure S7:**
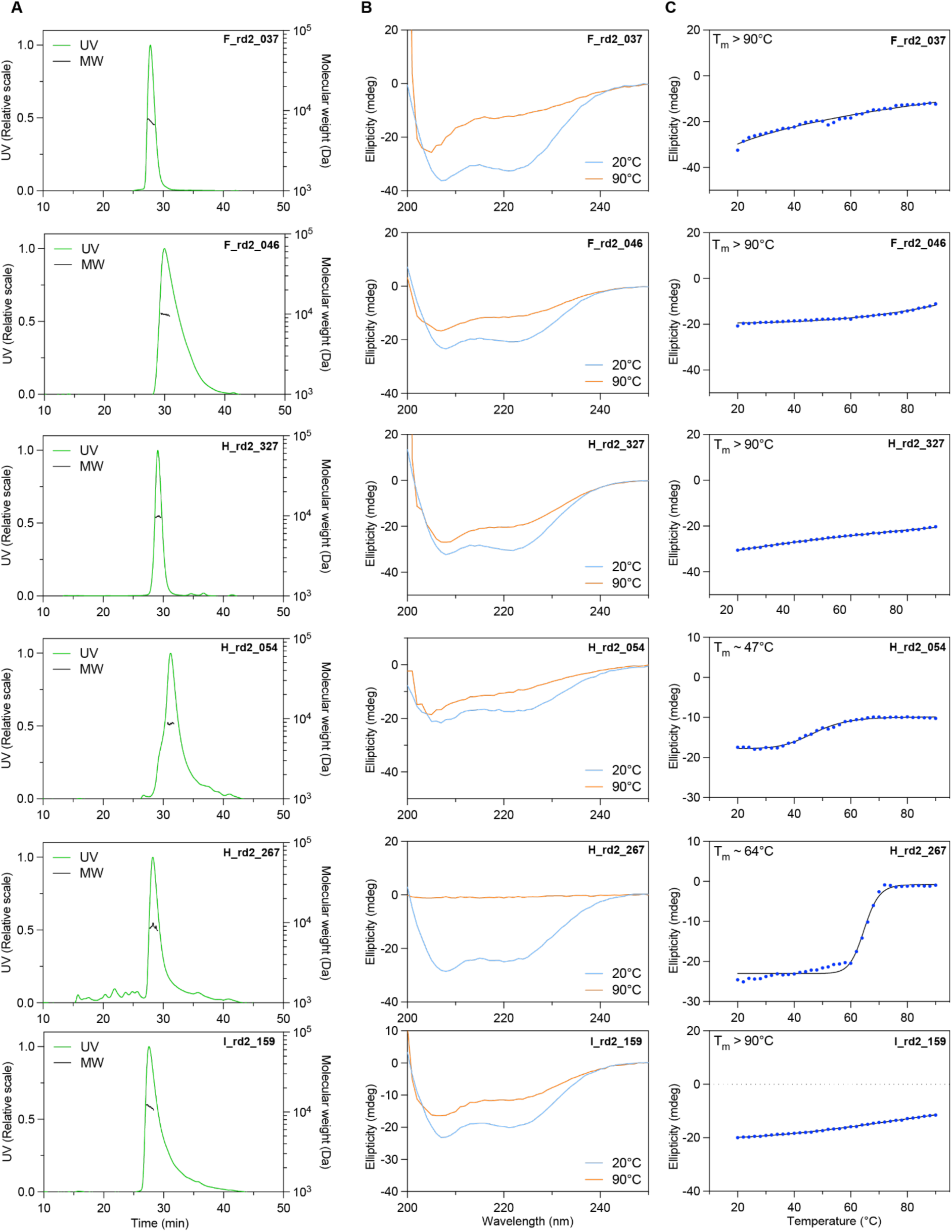
Biophysical characterization of purified binders (round 2). **A.** Oligomeric state determined by size-exclusion multi-angle light scattering (SEC-MALS). **B.** Protein folding of the purified binder measured by circular dichroism at 20°C (blue) or 90°C (orange). **C**. Thermal stability determined by measuring the ellipticity at 218 nm at increasing temperature.

**Supplementary Figure S8:**
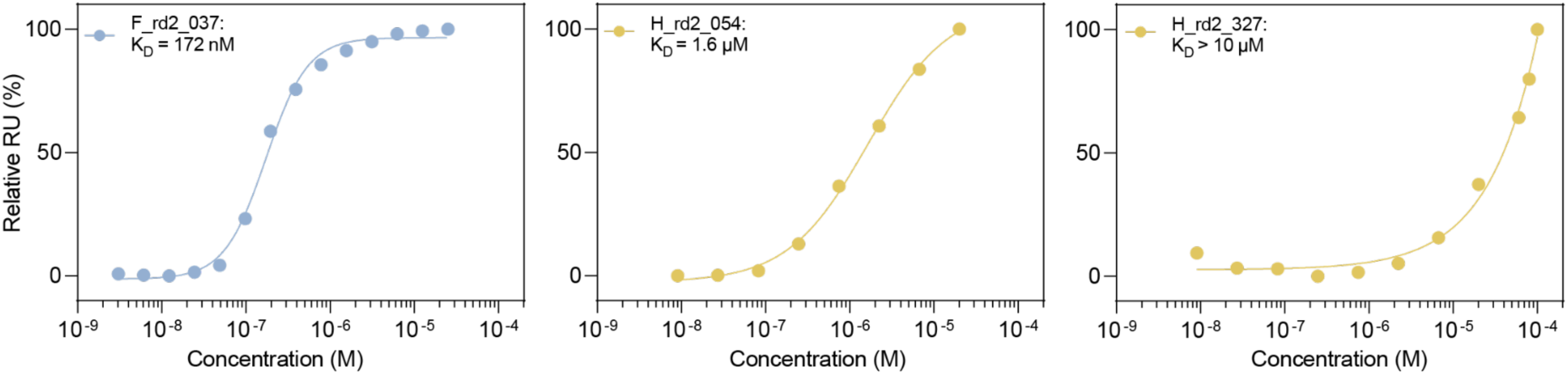
Surface plasmon resonance of selected rd2 binders. Affinity measurements of lower affinity rd2 binders performed by surface plasmon resonance (SPR). Steady state responses and dissociation constants were plotted using a nonlinear four-parameter curve fitting analysis.

**Supplementary Figure S9:**
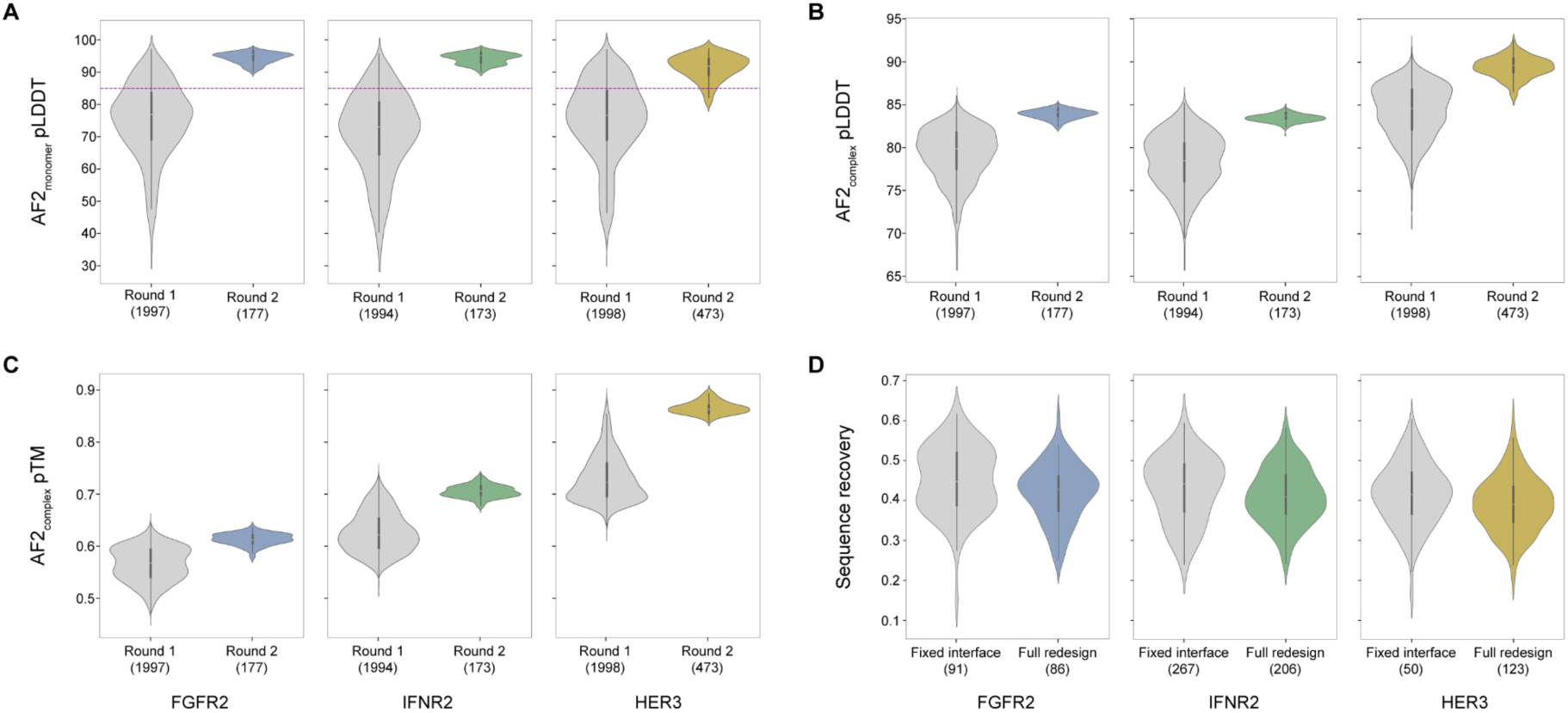
Computational metrics for round 2 optimized designs. Comparison of designs’ AF2 metrics before and after optimization in terms of **A.** Monomer pLDDT (dashed line set at pLDDT of 85%), **B.** Complex pLDDT, and **C.** Complex pTM. **D.** ProteinMPNN sequence recovery metric for round 2 designs after optimization with the two settings (i) Fixed interface and (ii) Full redesign. The results show that more than 50% of the round 1 binder sequence was redesigned in most cases.

**Supplementary Figure S10:**
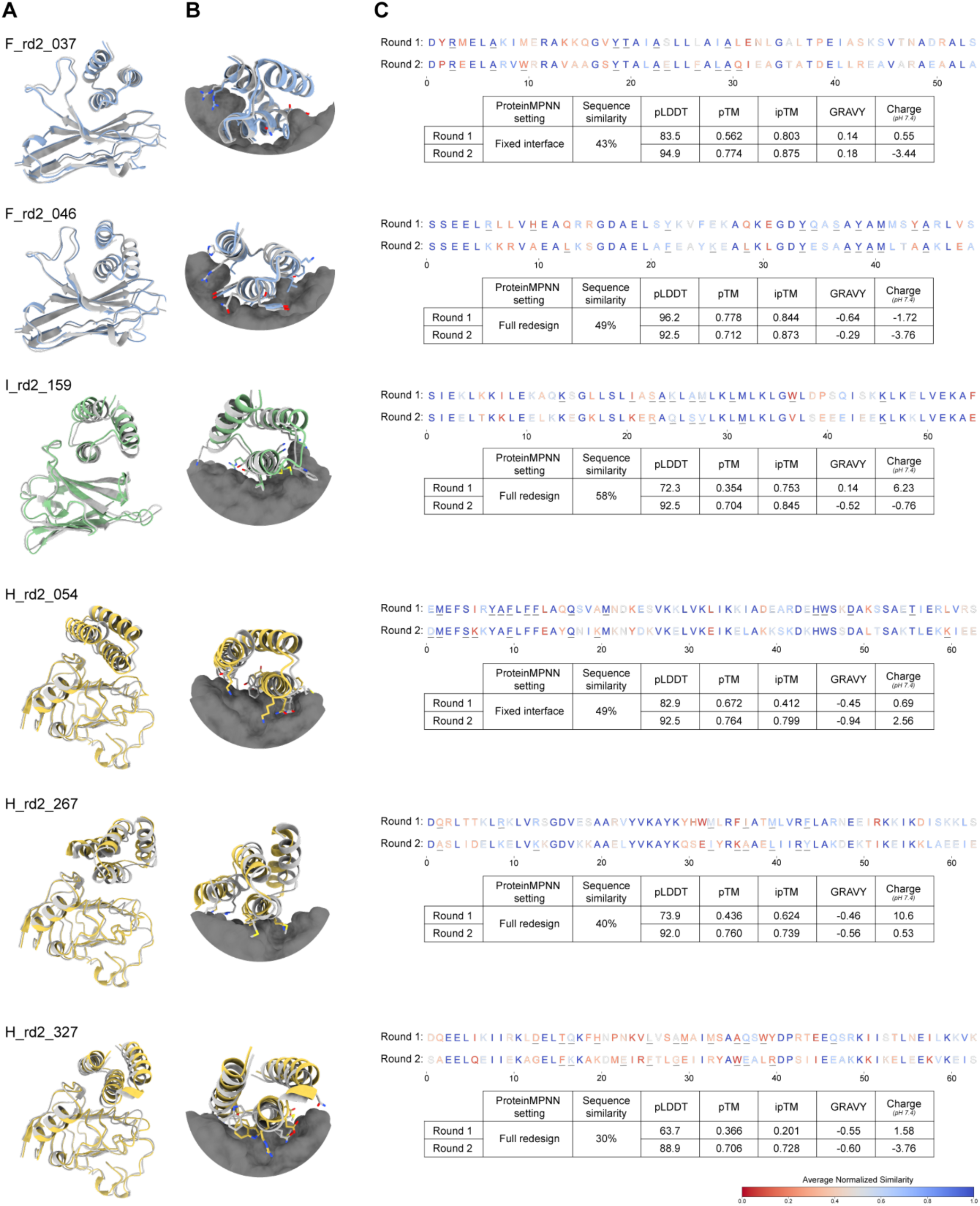
Computational analysis for round 2 experimentally validated designs. **A.** Structural alignment of round 2 designs’ AF2 multimer prediction (colored) to the Rosetta models (gray) of their originating designs from round 1. **B.** Close up on the designs’ interface with a selection of interface hotspots displayed in sticks. **C.** Design sequence alignment before and after optimization. The interface residues, 3.5Å from target interface, are underlined. The table shows a comparison of AF2 metrics and sequence properties including; AF2_monomer_ pLDDT, AF2_monomer_ pTM, AF2_complex_ ipTM, sequence similarity, GRAVY (Grand Average of Hydropathy) index and design charge at pH 7.4.

**Supplementary Figure S11:**
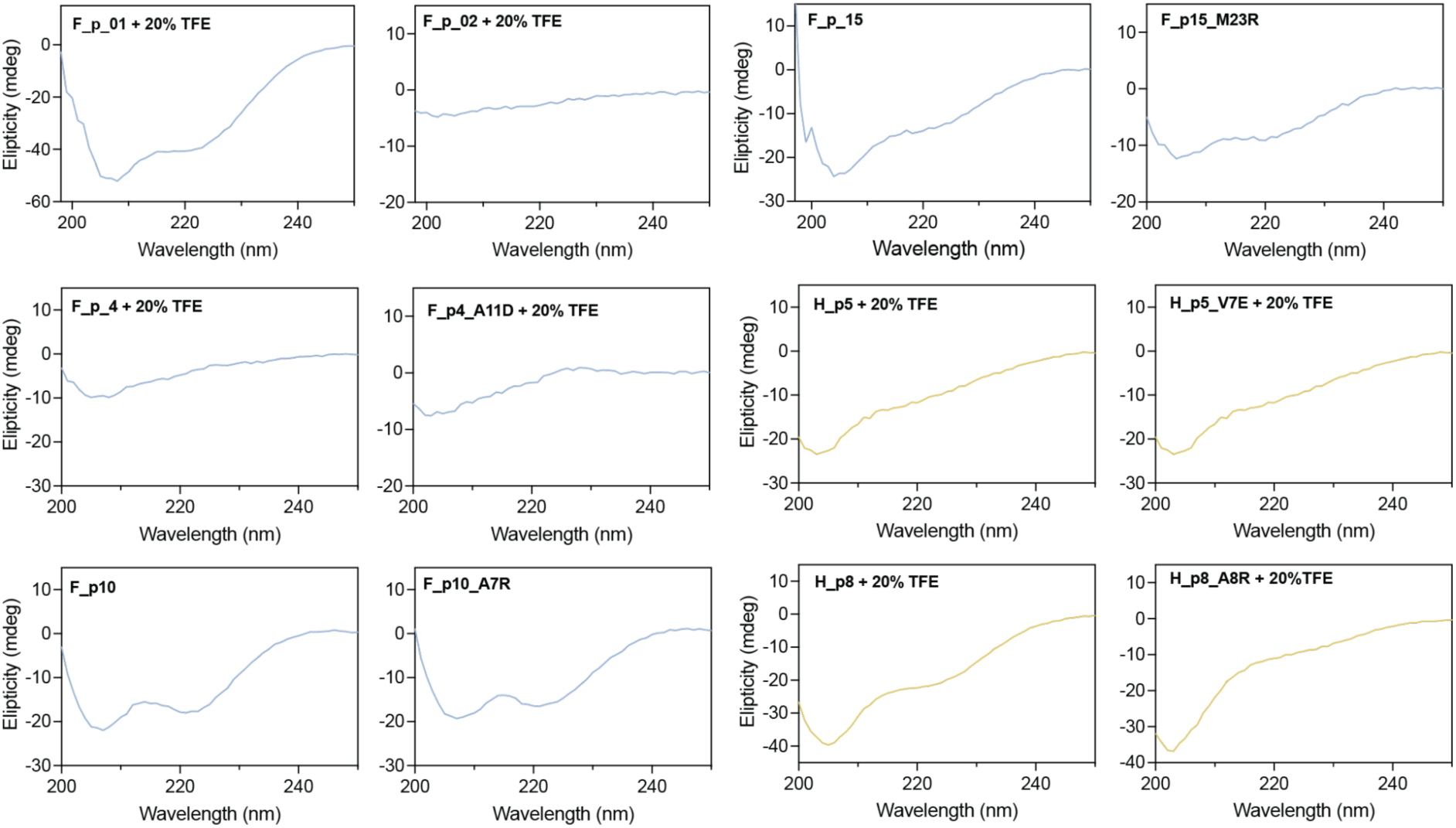
Circular dichroism measurement of target peptides and their respective knockout mutants. All peptides were measured at 100 µM, 20°C in PBS. Peptides F_01, F_02, F_p4, F_p4_A11D, H_p5, H_p5_V7E, H_p8 and H_p8_A8R were measured in the presence of 20% TFE.

**Supplementary Figure S12:**
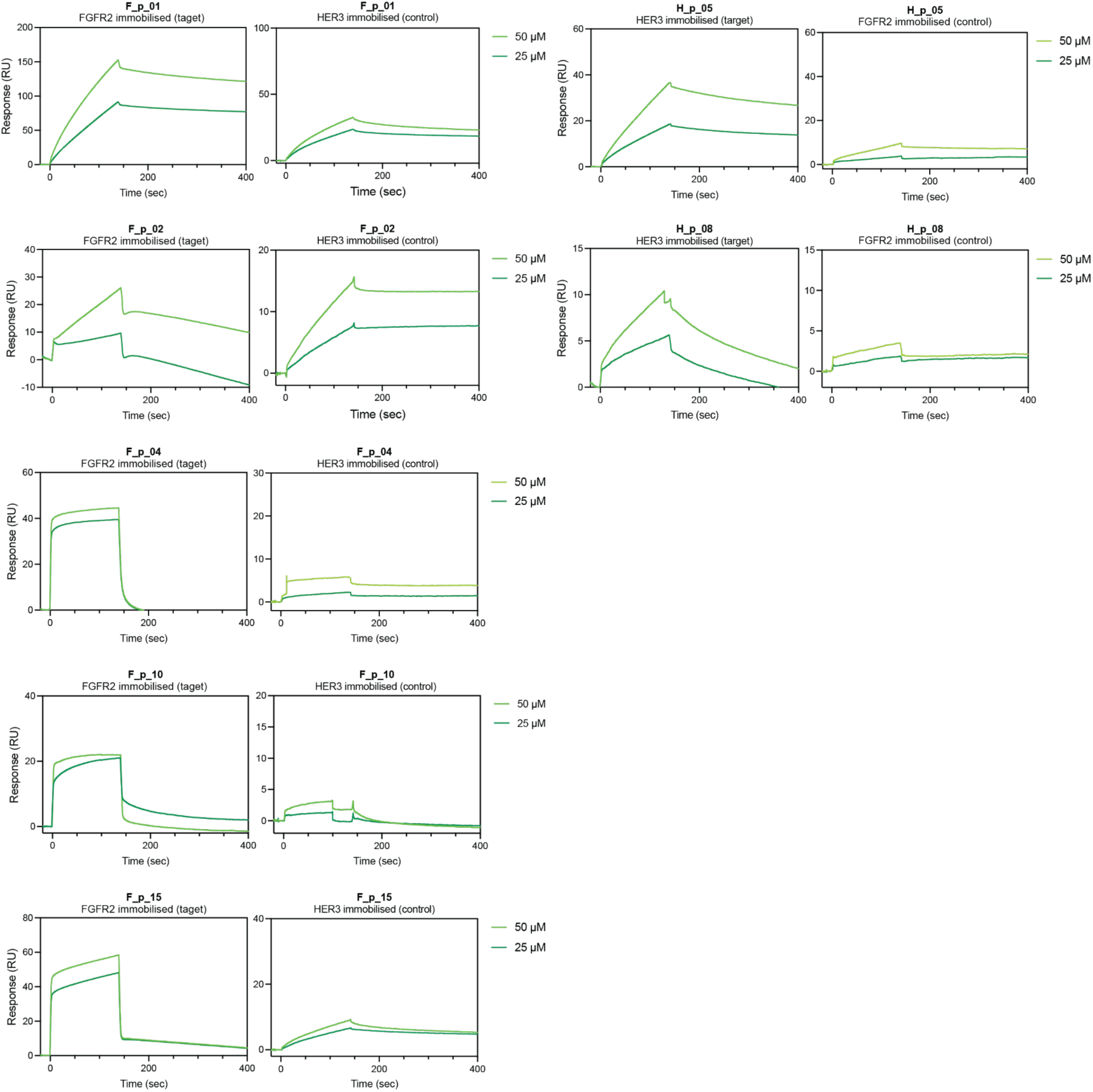
Surface plasmon resonance of selected peptides on target and unrelated protein to test for target specificity. Affinity measurements of successful peptides performed by surface plasmon resonance (SPR) against the target protein (left) and a control protein (right). All measurements were conducted with 50 µM and 25 µM of peptides. The ratio of immobilization levels of FGFR vs HER3 was 2:1, which is accounted for by the range of the y-axis. All measurements were conducted with 50 µM and 25 µM of peptides. Target specificity was determined by a lack of response of the peptide towards the control protein.

**Supplementary Table S1:** Summary of all sorted and tested designs. Designs were selected by yeast surface display based on their log-enrichment in binding versus non-binding populations. Selected designs were tested by yeast surface display for binding to the target protein with or without a point mutation at the interface or a competitor protein. Successful designs were then expressed (measured here in milligram of proteins per liter of bacteria culture) and characterized using various biophysical techniques (characterized designs are highlighted in bold). N/A: Not applicable. N/D: Not determined.

| Design | SS | Counts Non-Bind. | Counts Binding | Enrichment | Yeast binding | Point mutation specificity | Competition assay specificity | Expression [mg/L] | Characterized? |
| --- | --- | --- | --- | --- | --- | --- | --- | --- | --- |
| F_rd1_281 | S | 1279 | 20105 | 1.1964 | ++ | Yes | Yes | – | – |
| **F_rd1_702** | **H** | **132** | **10117** | **1.8845** | **+++** | **Yes** | **Yes** | **6.2** | **Yes** |
| **F_rd1_1013** | **H** | **4** | **18502** | **3.6652** | **+++** | **Yes** | **Yes** | **9.4** | **Yes** |
| F_rd1_1835 | H | 1152 | 113659 | 1.9942 | +++ | Yes | Yes | – | – |
| F_rd1_1837 | H | 217 | 99802 | 2.6627 | ++ | – | N/A | N/A | – |
| F_rd1_1854 | H | 40 | 30987 | 2.8891 | ++ | – | N/A | N/A | – |
| F_rd1_1893 | H | 207 | 45743 | 2.3444 | +++ | – | N/A | N/A | – |
| F_rd1_1929 | H | 168 | 14670 | 1.9411 | +++ | – | N/A | N/A | – |
| F_rd1_1933 | H | 4585 | 57672 | 1.0996 | +++ | – | N/A | N/A | – |
| F_rd1_1989 | H | 181 | 32188 | 2.25 | +++ | – | N/A | N/A | – |
| I_rd1_360 | S | 2920 | 224637 | 1.8861 | + | Yes | N/D | – | – |
| **H_rd1_309** | **S** | **1418** | **174633** | **2.0905** | **++** | **Yes** | **Yes** | **25.2** | **Yes** |
| H_rd1_1843 | H | 1179 | 342405 | 2.463 | ++ | – | N/A | N/A | – |
| H_rd1_1862 | H | 1953 | 30070 | 1.1874 | + | Yes | – | N/A | – |
| **F_rd2_037** | **H** | **42** | **23984** | **2.7567** | **+++** | **Yes** | **Yes** | **1.8** | **Yes** |
| **F_rd2_046** | **H** | **13** | **12680** | **2.9891** | **+++** | **Yes** | **Yes** | **11.3** | **Yes** |
| F_rd2_215 | H | 12 | 19132 | 3.2025 | +++ | N/A | N/A | – | – |
| **I_rd2_159** | **H** | **61018** | **1363868** | **1.3493** | **++** | **Yes** | **Yes** | **0.25** | **Yes** |
| H_rd2_054 | H | 85 | 656 | 0.8874 | + | Yes | – | 1.3 | Yes |
| **H_rd2_267** | **H** | **76** | **278579** | **3.5641** | **+++** | **Yes** | **Yes** | **0.57** | **Yes** |

**Supplementary Table S2:** Summary of round 2 designs’ *in silico* successes. Designs from round 1 were refined by applying a number of machine learning methods. First, the sequences were optimized using ProteinMPNN in two settings, (i) fh: fixing the interface residues within 3.5Å from the target and optimizing the scaffold sequence, and (ii) da: full re-design. For each design, 25 sequences were generated using each setting. Next, for each design, the best 5 unique sequences from each setting based on the global score were input into the AF2 monomer model to predict their *in silico* folding. The designs were filtered based on (i) pLDDT (≧90, except for HER3 designs: ≧80) and (ii) prediction C⍺ RMSD to model (≦1.5 Å). Then, the filtered designs’ interface was scored employing complex prediction using the AF2 monomer model. Finally, a selection of best designs with ipTM ≧0.7 was forwarded to experimental testing. Total: Percentage of total number of designs. AF2_m_: Percentage of AF2 monomer filtered designs.

|  | FGFR2 | HER3 | IFNAR2 |
| --- | --- | --- | --- |
| # Round 1 designs | 1'997 | 1'998 | 1'994 |
| Sequence optimization with ProteinMPNN | 19'860<br>(9'930/setting) | 19'980<br>(9'990/setting) | 19'940<br>(9'970/setting) |
| Filtering based on designs' foldability | 2'710<br>Total: 13.6% | 13'248<br>Total: 66.3% | 1'478<br>Total: 7.4% |
| Filtering based on complex interface quality | 2'235<br>Total: 11.3%<br>AF2<sub>m</sub>: 82.5% | 545<br>Total: 2.7%<br>AF2<sub>m</sub>: 4.1% | 581<br>Total: 2.9%<br>AF2<sub>m</sub>: 39.3% |
| Experimentally tested designs | 177 | 473 | 173 |

**Supplementary Table S3:**
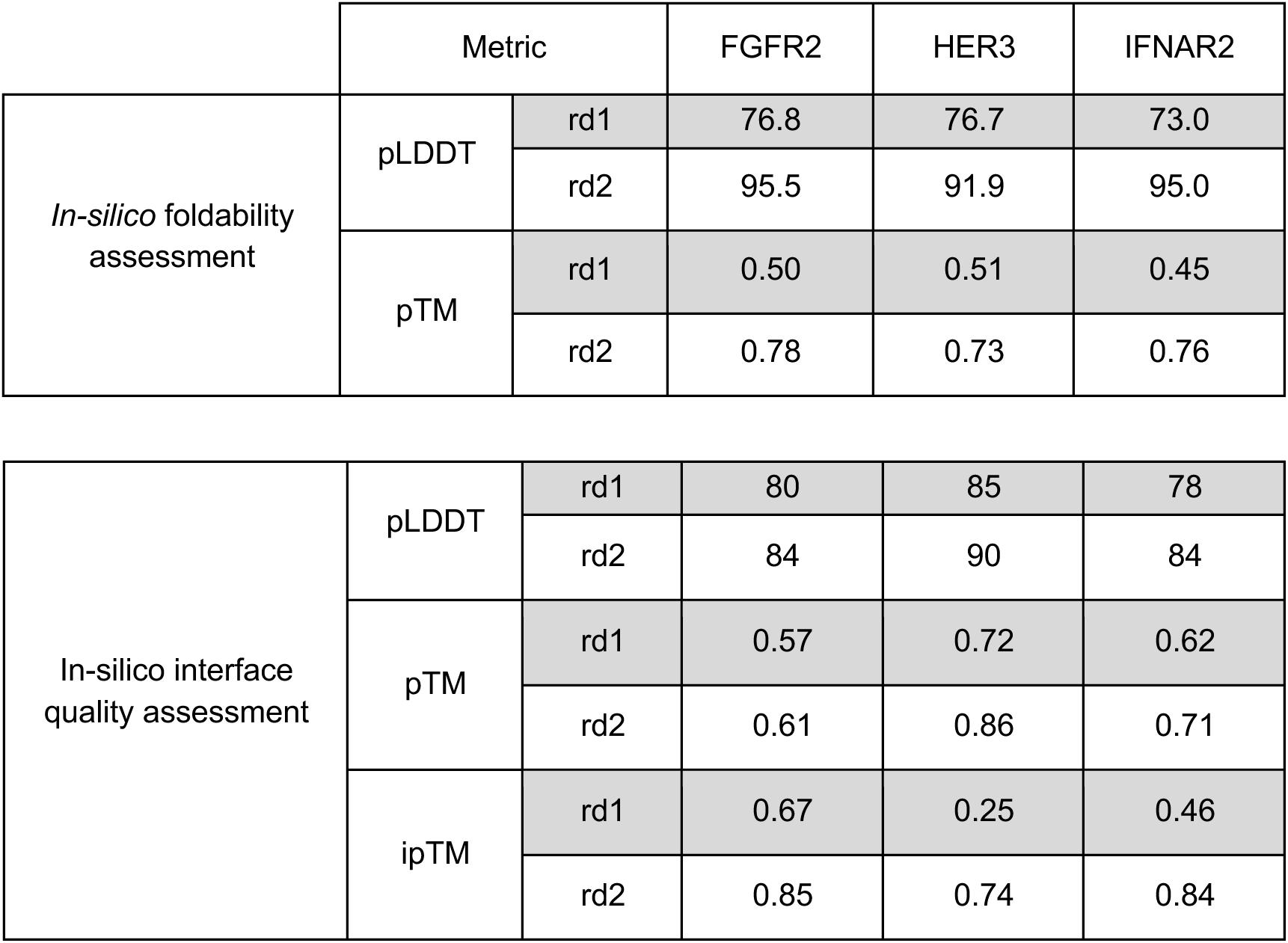
Comparison of AlphaFold2 metrics between rd1 and rd2 designs. Comparing the median values for the experimentally tested designs using yeast display surface from both design rounds. pLDDT: predicted local-distance difference test. pTM: The estimate of the TM-score. ipTM: interface pTM score

**Supplementary Table S4:**
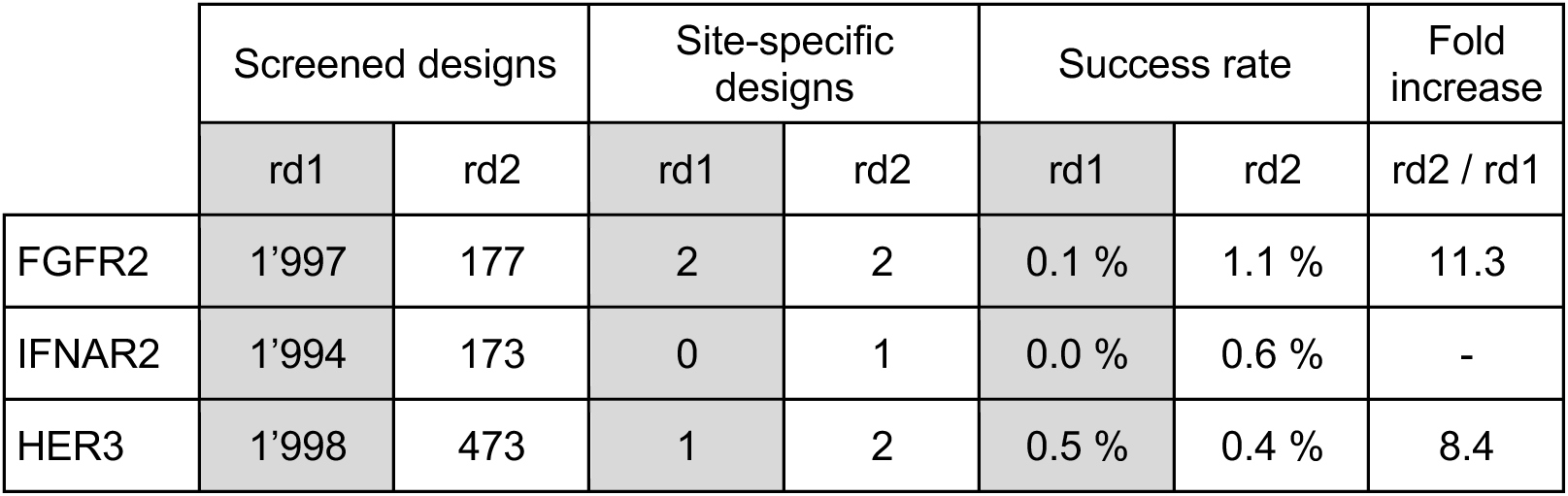
Comparison of experimental success rates between rd1 and rd2 designs. Comparing the number of screened designs using yeast surface display and the *in vitro* validated ones in terms of site-specificity and binding affinity.

|  | Screened designs |  | Site-specific designs |  | Success rate |  | Fold increase |
| --- | --- | --- | --- | --- | --- | --- | --- |
|  | rd1 | rd2 | rd1 | rd2 | rd1 | rd2 | rd2 / rd1 |
| FGFR2 | 1'997 | 177 | 2 | 2 | 0.1 % | 1.1 % | 11.3 |
| IFNAR2 | 1'994 | 173 | 0 | 1 | 0.0 % | 0.6 % | - |
| HER3 | 1'998 | 473 | 1 | 2 | 0.5 % | 0.4 % | 8.4 |

**Supplementary Table S5:**
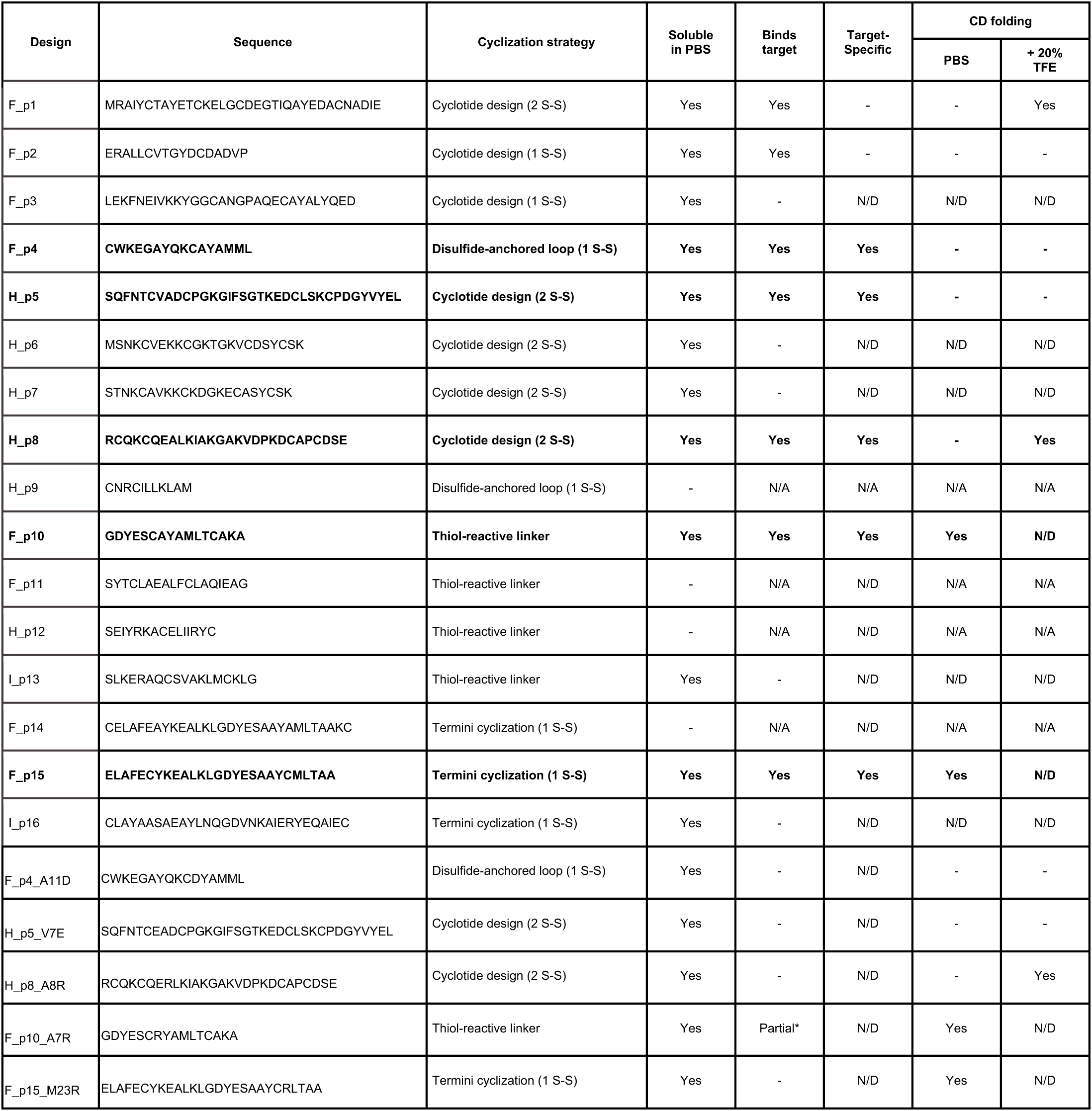
Summary of all tested peptides’ experimental characterization. Peptides generated from the extracted binding motifs of rd1 and rd2 mini-protein binder hits, which were soluble in PBS, were tested using SPR to evaluate their binding to their respective target proteins (reported as: Binds target). Target specificity was further assessed using SPR by measuring the peptides’ affinity for a control protein (reported as: Target-specificity) and evaluating the binding of peptide interface knockout mutants to the desired target (target-specific designs are highlighted in bold). Cyclization strategies:

- Thiol-reactive linker: binding helical motifs were stabilized by introducing two cysteine residues, enabling the incorporation of thiol-reactive linker (i.e., 4,4’-Bis(bromomethyl)biphenyl).
- Termini cyclization: binding motifs were cyclized by introducing a cysteine residue at each terminus to facilitate disulfide bridge formation.
- Disulfide-anchored loop: binding motifs were cyclized by adding of a loop anchored to the motif via disulfide bond, with cysteine residues positioned at each terminus.
- Cyclotide design: binding motifs with disulfide-anchored loop were stabilized further by incorporating them into cyclized peptides of up to 38 residues using RFDiffusion. To further enhance stability, Rosetta was employed to introduce a second, internal disulfide bond when suitable. N/A: Not applicable. N/D: Not determined. CD: Circular Dichroism. The number of disulfide bonds engineered into the peptide (S-S). *Partial: F_p10_A7R shows binding to FGFR2, albeit lesser than F_p10.

**Supplementary Table S6:**
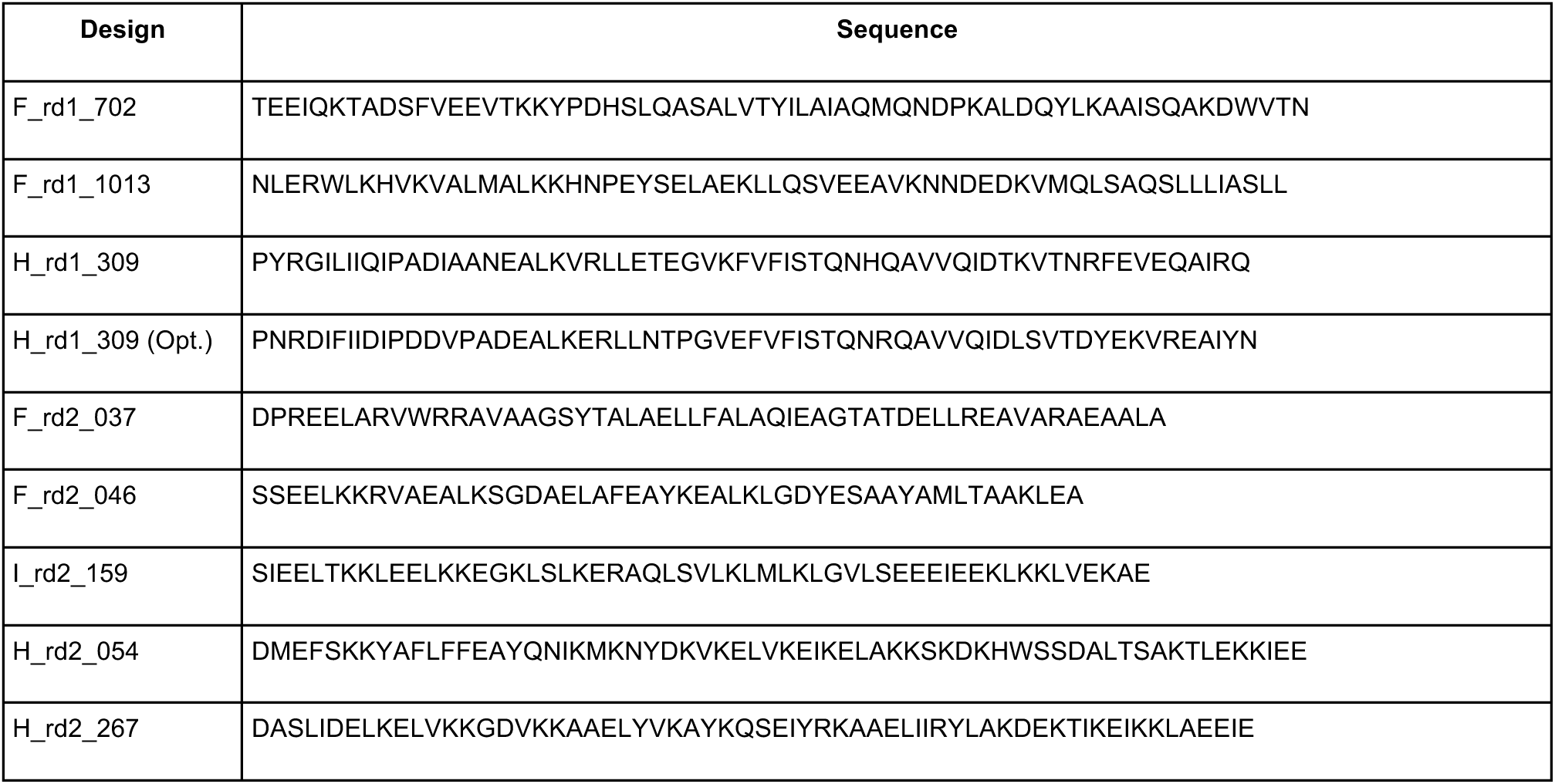
Expressed protein sequences from rd1 and rd2 design rounds.

## References

[1] D. Bausch-Fluck, E. S. Milani, and B. Wollscheid, “Surfaceome nanoscale organization and extracellular interaction networks,” Curr. Opin. Chem. Biol., vol. 48, pp. 26–33, Feb. 2019, doi: 10.1016/j.cbpa.2018.09.020.

[2] D. Bausch-Fluck et al., “The in silico human surfaceome,” Proc. Natl. Acad. Sci., vol. 115, no. 46, pp. E10988–E10997, 2018, doi: 10.1073/pnas.1808790115.

[3] C. M. Bryan et al., “Computational design of a synthetic PD-1 agonist,” Proc. Natl. Acad. Sci., vol. 118, no. 29, p. e2102164118, Jul. 2021, doi: 10.1073/pnas.2102164118.

[4] S. M. Swain, M. Shastry, and E. Hamilton, “Targeting HER2-positive breast cancer: advances and future directions,” Nat. Rev. Drug Discov., vol. 22, no. 2, pp. 101–126, Feb. 2023, doi: 10.1038/s41573-022-00579-0.

[5] L. Cao et al., “Design of protein-binding proteins from the target structure alone,” Nature, vol. 605, no. 7910, pp. 551–560, 2022.

[6] H. Khakzad, I. Igashov, A. Schneuing, C. Goverde, M. Bronstein, and B. Correia, “A new age in protein design empowered by deep learning,” Cell Syst., vol. 14, no. 11, pp. 925– 939, Nov. 2023, doi: 10.1016/j.cels.2023.10.006.

[7] R. Krishna et al., “Generalized biomolecular modeling and design with RoseTTAFold All-Atom,” Science, vol. 384, no. 6693, p. eadl2528, Apr. 2024, doi: 10.1126/science.adl2528.

[8] N. Bennett et al., “Improving de novo protein binder design with deep learning,” bioRxiv, 2022.

[9] J. L. Watson et al., “De novo design of protein structure and function with RFdiffusion,” Nature, vol. 620, no. 7976, pp. 1089–1100, Aug. 2023, doi: 10.1038/s41586-023-06415-8.

[10] P. Gainza et al., “De novo design of protein interactions with learned surface fingerprints,” Nature, pp. 1–9, 2023.

[11] J. Jumper et al., “Highly accurate protein structure prediction with AlphaFold,” Nature, vol. 596, no. 7873, pp. 583–589, 2021.

[12] P. Gainza et al., “Deciphering interaction fingerprints from protein molecular surfaces using geometric deep learning,” Nat. Methods, vol. 17, no. 2, pp. 184–192, 2020.

[13] J. Kyte and R. F. Doolittle, “A simple method for displaying the hydropathic character of a protein,” J. Mol. Biol., vol. 157, no. 1, pp. 105–132, May 1982, doi: 10.1016/0022-2836(82)90515-0.

[14] J. K. Leman et al., “Macromolecular modeling and design in Rosetta: recent methods and frameworks,” Nat. Methods, vol. 17, no. 7, pp. 665–680, 2020.

[15] J. Dauparas et al., “Robust deep learning–based protein sequence design using ProteinMPNN,” Science, p. eadd2187, 2022.

[16] R. Evans et al., “Protein complex prediction with AlphaFold-Multimer,” Oct. 04, 2021. doi: 10.1101/2021.10.04.463034.

[17] K. H. Sumida et al., “Improving Protein Expression, Stability, and Function with ProteinMPNN,” J. Am. Chem. Soc., vol. 146, no. 3, pp. 2054–2061, Jan. 2024, doi: 10.1021/jacs.3c10941.

[18] R. Yin, B. Y. Feng, A. Varshney, and B. G. Pierce, “Benchmarking AlphaFold for protein complex modeling reveals accuracy determinants,” Protein Sci., vol. 31, no. 8, p. e4379, Aug. 2022, doi: 10.1002/pro.4379.

[19] S. A. Rettie et al., “Cyclic peptide structure prediction and design using AlphaFold,” Feb. 26, 2023. doi: 10.1101/2023.02.25.529956.

[20] M. Pacesa et al., “BindCraft: one-shot design of functional protein binders,” Oct. 01, 2024. doi: 10.1101/2024.09.30.615802.

[21] S. Vázquez Torres et al., “De novo design of high-affinity binders of bioactive helical peptides,” Nature, vol. 626, no. 7998, pp. 435–442, Feb. 2024, doi: 10.1038/s41586-023-06953-1.

[22] S. J. Fleishman et al., “RosettaScripts: A Scripting Language Interface to the Rosetta Macromolecular Modeling Suite,” PLoS ONE, vol. 6, no. 6, p. e20161, Jun. 2011, doi: 10.1371/journal.pone.0020161.

[23] B. Coventry and D. Baker, “Protein sequence optimization with a pairwise decomposable penalty for buried unsatisfied hydrogen bonds,” PLOS Comput. Biol., vol. 17, no. 3, p. e1008061, Mar. 2021, doi: 10.1371/journal.pcbi.1008061.

[24] D.-A. Silva, B. E. Correia, and E. Procko, “Motif-Driven Design of Protein–Protein Interfaces,” in Computational Design of Ligand Binding Proteins, vol. 1414, B. L. Stoddard, Ed., in Methods in Molecular Biology, vol. 1414., New York, NY: Springer New York, 2016, pp. 285–304. doi: 10.1007/978-1-4939-3569-7_17.

[25] H. M. Berman, “The Protein Data Bank,” Nucleic Acids Res., vol. 28, no. 1, pp. 235–242, Jan. 2000, doi: 10.1093/nar/28.1.235.

[26] A. R. Tobin, R. Crow, D. V. Urusova, J. C. Klima, N. H. Tolia, and E. Strauch, “Inhibition of a malaria host–pathogen interaction by a computationally designed inhibitor,” Protein Sci., vol. 32, no. 1, p. e4507, Jan. 2023, doi: 10.1002/pro.4507.

[27] G. Bhardwaj et al., “Accurate de novo design of hyperstable constrained peptides,” Nature, vol. 538, no. 7625, pp. 329–335, Oct. 2016, doi: 10.1038/nature19791.

[28] T. W. Linsky et al., “Sampling of structure and sequence space of small protein folds,” Nat. Commun., vol. 13, no. 1, p. 7151, Nov. 2022, doi: 10.1038/s41467-022-34937-8.

[29] G. J. Rocklin et al., “Global analysis of protein folding using massively parallel design, synthesis, and testing,” Science, vol. 357, no. 6347, pp. 168–175, Jul. 2017, doi: 10.1126/science.aan0693.

[30] P. J. A. Cock et al., “Biopython: freely available Python tools for computational molecular biology and bioinformatics,” Bioinformatics, vol. 25, no. 11, pp. 1422–1423, Jun. 2009, doi: 10.1093/bioinformatics/btp163.

[31] P. Kunzmann and K. Hamacher, “Biotite: a unifying open source computational biology framework in Python,” BMC Bioinformatics, vol. 19, no. 1, p. 346, Dec. 2018, doi: 10.1186/s12859-018-2367-z.

[32] P. Kunzmann et al., “Biotite: new tools for a versatile Python bioinformatics library,” BMC Bioinformatics, vol. 24, no. 1, p. 236, Jun. 2023, doi: 10.1186/s12859-023-05345-6.

[33] C. A. Goverde et al., “Computational design of soluble and functional membrane protein analogues,” Nature, vol. 631, no. 8020, pp. 449–458, Jul. 2024, doi: 10.1038/s41586-024-07601-y.

[34] M. Mirdita, K. Schütze, Y. Moriwaki, L. Heo, S. Ovchinnikov, and M. Steinegger, “ColabFold: making protein folding accessible to all,” Nat. Methods, vol. 19, no. 6, pp. 679–682, 2022.

[35] G. Chao, W. L. Lau, B. J. Hackel, S. L. Sazinsky, S. M. Lippow, and K. D. Wittrup, “Isolating and engineering human antibodies using yeast surface display,” Nat. Protoc., vol. 1, no. 2, pp. 755–768, Aug. 2006, doi: 10.1038/nprot.2006.94.

[36] P. Diderich et al., “Phage Selection of Chemically Stabilized α-Helical Peptide Ligands,” ACS Chem. Biol., vol. 11, no. 5, pp. 1422–1427, May 2016, doi: 10.1021/acschembio.5b00963.

[37] P. Hosseinzadeh et al., “Comprehensive computational design of ordered peptide macrocycles,” Science, vol. 358, no. 6369, pp. 1461–1466, Dec. 2017, doi: 10.1126/science.aap7577.

[38] P. J. Salveson et al., “Expansive discovery of chemically diverse structured macrocyclic oligoamides,” Science, vol. 384, no. 6694, pp. 420–428, Apr. 2024, doi: 10.1126/science.adk1687.

[39] T. Kosugi and M. Ohue, “Design of Cyclic Peptides Targeting Protein–Protein Interactions Using AlphaFold,” Int. J. Mol. Sci., vol. 24, no. 17, p. 13257, Aug. 2023, doi: 10.3390/ijms241713257.

[40] M. J. Abraham et al., “GROMACS: High performance molecular simulations through multi-level parallelism from laptops to supercomputers,” SoftwareX, vol. 1–2, pp. 19–25, Sep. 2015, doi: 10.1016/j.softx.2015.06.001.

[41] H. Geng, F. Jiang, and Y.-D. Wu, “Accurate Structure Prediction and Conformational Analysis of Cyclic Peptides with Residue-Specific Force Fields,” J. Phys. Chem. Lett., vol. 7, no. 10, pp. 1805–1810, May 2016, doi: 10.1021/acs.jpclett.6b00452.

[42] S. Miyamoto and P. A. Kollman, “Settle: An analytical version of the SHAKE and RATTLE algorithm for rigid water models,” J. Comput. Chem., vol. 13, no. 8, pp. 952–962, Oct. 1992, doi: 10.1002/jcc.540130805.

[43] B. Hess, H. Bekker, H. J. C. Berendsen, and J. G. E. M. Fraaije, “LINCS: A linear constraint solver for molecular simulations,” J. Comput. Chem., vol. 18, no. 12, pp. 1463–1472, Sep. 1997, doi: 10.1002/(SICI)1096-987X(199709)18:12<1463::AID-JCC4>3.0.CO;2-H.

[44] M. Parrinello and A. Rahman, “Polymorphic transitions in single crystals: A new molecular dynamics method,” J. Appl. Phys., vol. 52, no. 12, pp. 7182–7190, Dec. 1981, doi: 10.1063/1.328693.

